# A non-cell autonomous actin redistribution enables isotropic retinal growth

**DOI:** 10.1101/293951

**Authors:** Marija Matejčić, Guillaume Salbreux, Caren Norden

**Affiliations:** Max Planck Institute of Molecular Cell Biology and Genetics, Pfotenhauerstraße 108, 01307 Dresden, Germany; The Francis Crick Institute 1 Midland Road, London, NW1, UK

**Author notes:** Corresponding Authors: Caren Norden (C.N.) Guillaume Salbreux (G.S.).

## Abstract

Tissue shape is often established early in development and needs to be scaled isotropically during growth. However, the cellular contributors and ways in which cells interact inside tissues to enable coordinated isotropic tissue scaling are not yet understood. Here, we follow cell and tissue shape changes in the zebrafish retinal neuroepithelium, which forms a cup with a smooth surface early in development and maintains this architecture as it grows. By combining 3D analysis and theory, we show that a global increase in cell height is necessary to maintain this tissue shape during growth. Timely cell height increase is governed by non-cell autonomous actin redistribution. Blocking actin redistribution and cell height increase perturbs isotropic scaling and leads to disturbed, folded tissue shape. Taken together, our data show how global changes in cell shape enable isotropic growth of the developing retinal neuroepithelium, a concept that could also apply to other systems.

## Introduction

Acquiring the correct size and shape during development is a crucial prerequisite for tissue and organ function. In many tissues, such as the drosophila salivary gland [1] or the vertebrate retina [2], shape characteristics are established early in development and need to be retained throughout growth necessitating an isotropic rescaling of the initial tissue shape (Fig 1A). How such a uniform, isotropic rescaling is achieved through cell and tissue level processes is not yet well explored.

**Figure 1.**
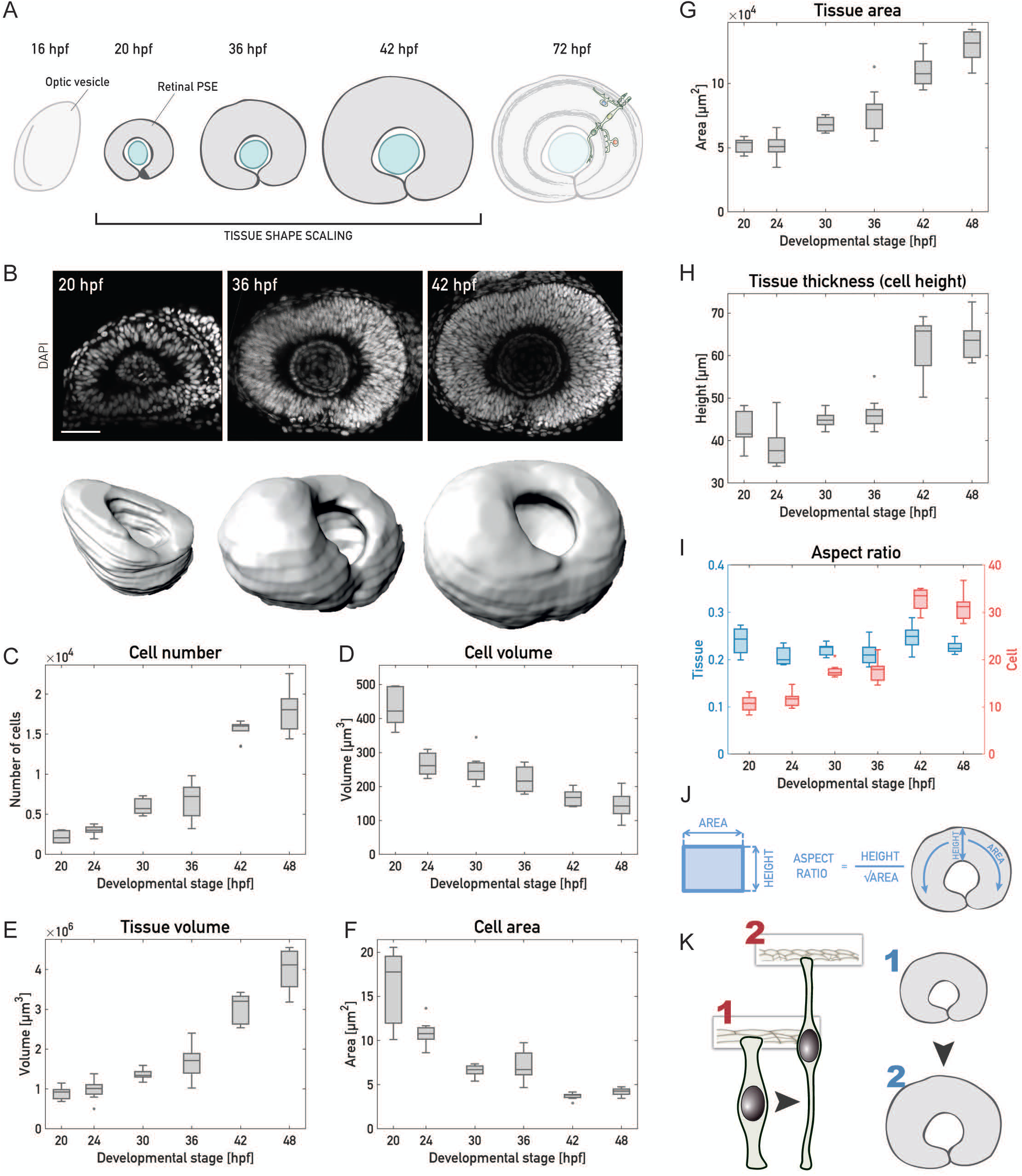
Changes of cell shapes allow for tissue shape maintenance during retinal PSE growth. **(A)**, Schematic of retinal development. After the optic vesicle transforms into the optic, cells in the retinal PSE proliferate and the tissue maintains its shape (20-42 hpf) to ultimately give rise to the laminated neuronal retina. **(B)**, Upper panels: Representative images of the retinal PSE in the proliferative growth phases. Nuclei are labelled with DAPI. Scale bar: 50 µm. Lower panels: 3D surfaces used in the retinal PSE tissue-wide growth analysis at 20 hpf, 36 hpf and 42 hpf. Surfaces were manually segmented, rendered and analyzed using Imaris 8.3 (Bitplane). Related to S1 Movie. **(C)**, Total cell number increase during retinal PSE growth. **(D)**, Cell volume decrease, as measured and corrected from mitotic cell volumes. **(E)**, Tissue volume increase from manually segmented retinal tissues. **(F)**, Mean cell area decrease. Cell area is an average of cell apical and basal areas, calculated from cell number and tissue areas. **(G)**, Tissue area increase. Area shown is an average of the apical and basal tissue area. **(H)**, Tissue thickness increase as measured by cell height. **(I)**, Tissue (blue) and cell (red) aspect ratios. **(J)**, Aspect ratios were calculated by dividing tissue height by the square root of the average tissue area (plotted in **G**), or average cell area (**F**). **(K)**, Schematic representation of the shape changes during retinal PSE growth. PSE cells change their shape by elongating and becoming thinner, while the overall tissue shape remains. (**C-I**) 10 samples/stage. On the boxplots, the central mark marks the median, the bottom and top box limits indicate the 25th and 75th percentile, respectively, the whiskers indicate the most extreme data points that are not considered outliers, outliers are plotted as points.

Some of the molecular factors governing cell and tissue geometry have been extensively studied. Both microtubule reorganization and cortical actomyosin rearrangements can influence shapes of individual epithelial cells, as well as the shape of whole tissues [3–10]. When studying these factors however, most investigations so far focused on anisotropic tissue shape changes occurring during development, while isotropic growth and shape maintenance is not as well understood. In addition, most studies explored tissue growth and shape in two dimensions [5,9,11–13]. Thus, a thorough evaluation of how tissue growth is coordinated by cell and tissue parameters in 3D is currently lacking.

One tissue that allows to analyze cell and tissue shape during growth is the curved vertebrate retinal neuroepithelium. The retinal neuroepithelium establishes its shape early in development by formation of the optic cup [2,14] and grows continuously afterwards (Fig 1A). During this proliferative growth, the neuroepithelium is pseudostratified [15], with cells attached to both the apical side and the basal lamina. Nuclei are located along the apicobasal axis, except in mitosis, as cells divide apically [16]. Such pseudostratified epithelia (PSE) are evolutionary conserved and serve as organ precursors in a wide range of species [15]. In the retinal neuroepithelium as well as in most other PSE, cells progressively increase their height as the tissue proliferates and grows [15,17–19]. So far, it is not understood what drives this cell elongation over development and whether and how it links to the overall tissue growth.

To investigate how growth and shape coordinate in the developing retinal neuroepithelium, we used zebrafish to analyze cell and tissue wide shape changes during proliferative growth. We obtained a quantitative tissue-wide 3D dataset, which allowed us to break down 3D growth into individual cellular components through balance equations. Our analysis revealed that isotropic growth relies on apico-basal cell elongation. This cell elongation is governed by cellular actin redistribution that occurs in a non-cell autonomous manner. Furthermore, cell elongation occurs in concert with continued proliferation and needs an unperturbed extracellular matrix (ECM). Our data is validated by a simplified theoretical model and balance equations that account for the main aspects of tissue changes. If actin redistribution and thereby cell height increase is impaired, tissue shape maintenance does not take place and the tissue gets distorted upon further growth.

## Results

### During retinal PSE growth, single cell aspect ratios change while the overall tissue aspect ratio is maintained

In order to understand how cell and tissue shape changes are linked during growth of the proliferating retinal PSE, we quantified cell and tissue geometry of the retinal neuroepithelium over development. By performing tissue-wide 3D analysis (Fig 1B, S1 Movie), we analyzed growth in 4-6 hour intervals between 20 hours post fertilization (hpf), when morphogenesis of the optic cup is complete (Fig 1A, B), and 48 hpf, a stage when neuronal differentiation is ongoing, (Fig 1A, B, 10 embryos/time-point). While apoptosis was negligible during this phase (S1 Fig)[20], the total cell number increased 8-fold, from ~2,200 to ~18,000 (Fig 1C). Concurrently, single cell volumes decreased by almost two thirds, from 440 to 150 µm^3^ (Fig 1D). Overall, the tissue volume increased exponentially by a factor 3.4 over 28 hours, corresponding to a rate of ~0.055/h and a volume doubling time of 12.5 hours (Fig 1E).

In addition to decreasing their volume, cells changed their shape, becoming thinner (Fig 1F) and more apico-basally elongated (Fig 1H). The largest change in aspect ratio and thereby cell shape occurred between 36 hpf and 42 hpf (Fig 1I), as did the largest increase in cell numbers (Fig 1C), suggesting differences between an early (before 36 hpf) and later growth phase (after 36 hpf). We next examined the overall changes in tissue area (Fig 1G) and height (Fig 1H) over time (Fig 2A). Before 36 hpf, tissue height stayed fairly constant and most of the volume growth was accounted for by area expansion. After 36 hpf however, a significant increase of tissue height was seen, which accounted for a larger fraction of tissue growth. While single cells elongated and changed their aspect ratio (Fig 1I), the tissue area (Fig 1G) and cell height (Fig 1H) roughly scaled with each other, such that the aspect ratio of the tissue remained constant (Fig 1I-K). This may seem in contradiction with the observation that area expansion occurs at constant height before 36 hpf. However, during this phase, the area expansion is small (Fig 1G) and the tissue aspect ratio is therefore not significantly affected (Fig 1I).

**Figure 2.**
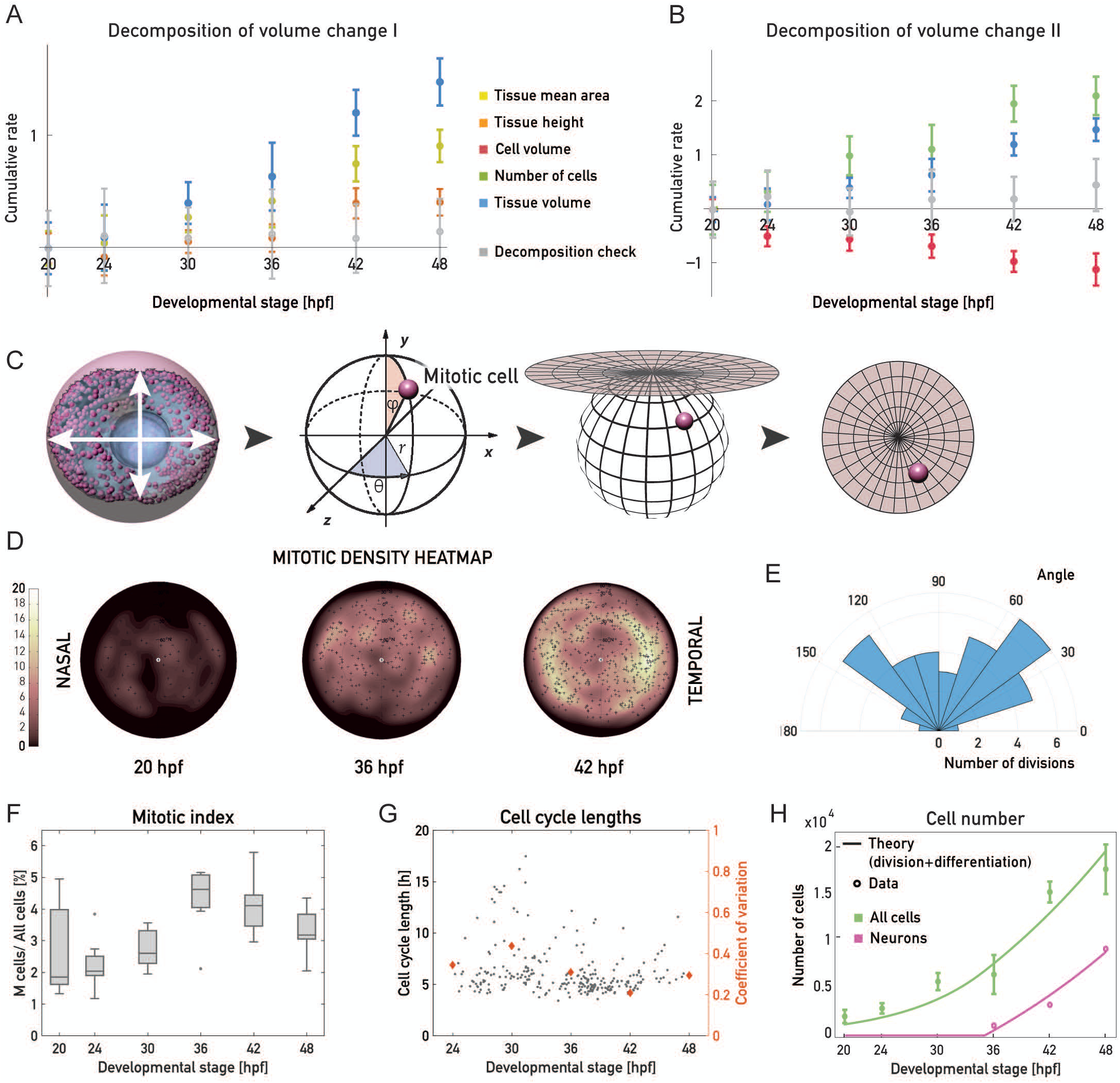
Proliferative growth of the retinal PSE is isotropic and homogeneous. **(A)**, Cumulative rate of change for the number of cells (green), tissue height (orange), and tissue average area (yellow). Grey data points correspond to the sum of these three independently measured rates of changes, which add up to 0 within experimental standard deviation. **(B),** Cumulative rate of change for the tissue volume (orange), number of cells (green) and cell volume (blue). Grey data points correspond to the sum of these three independently measured rates of changes, which should add up to 0 within experimental standard deviation. **(C)**, Illustration of the mitotic distribution workflow to generate 2D mitotic density heatmaps. 3D Cartesian coordinates of every mitotic cell (see Methods) were transformed into polar, spherical coordinates, which were then projected into two dimensions using a density-preserving azimuthal projection tool. **(D)**, Typical 2D heatmaps of mitotic densities at 20, 36 and 42 hpf, results of transformations in **(C)**. 10 samples/stage. **(E)**, Rose plot of division angles, analyzed from a movie ~40 hpf (see S2 Movie). **(F)**, The mitotic index was calculated as the fraction of mitotic cells of all retinal PSE cells. This analysis was as well done by analyzing tissue-wide images in 3D. **(G)**, Cell cycle lengths of progenitor retinal PSE cells, analyzed by manual tracking of 254 cells from 20 embryos. Coefficient of variation (CoV) for each stage is plotted as orange diamonds (right y-axis). Data for CoV was binned as stage±3h (for example see S3 Movie). **(H)**, Total number of cells in the retina (green points, as in Fig 1D) and number of committed progenitors/neurons (magenta points), from data in S2 Fig Green and magenta lines: theoretical cell and neuron number, assuming a constant rate of division, and a 35% probability of dividing progenitors to produce 2 committed progenitors/neurons after 35 hpf (see Supplementary Notes). Data points are plotted as mean±SD.

Overall our measurements indicate that height and area expansion of the retinal PSE occur concurrently to maintain a fixed tissue aspect ratio.

### Retinal PSE growth is isotropic and homogeneous

The fact that the retinal PSE grows with a fixed aspect ratio suggested that mitotic events are distributed homogenously and isotropically in the plane of the epithelium. To test this notion, we analyzed the distribution of apical mitosis throughout the proliferative phase using a custom tool (Fig 2C and Materials and Methods) that allowed us to project the 3D coordinates of all apically dividing cells onto a 2D density heatmap (Fig 2D). This confirmed that mitotic cells were distributed evenly across the apical surface at all developmental stages (Fig 2D), making proliferative growth homogeneous. Isotropy of cell division angles in the plane of the epithelium was validated by live-imaging (Fig 2E and S2 Movie).

Our mitotic density heatmaps (Fig 2D) in combination with the mitotic index analysis (Fig 2F) showed that mitotic activity peaked at ~36-42 hpf. To understand whether this mitotic peak was due to an increased mitotic duration or due to overall cell cycle shortening, we analyzed cell cycle length by live-imaging of mosaically expressed nuclear marker, using light sheet microscopy to reduce phototoxicity (S3 Movie)[21]. Manual cell tracking (n=250, 20 embryos), revealed that as the length of mitosis remained constant (S2 Fig), average cell cycle lengths shortened from 7.3 h at 30 hpf, to 5.3 h at 42 hpf (Fig 2G), and became less variable, as seen by the reduced coefficient of variation (Fig 2G). Thus, the observed peak in proliferation between 36 hpf and 42 hpf resulted from an overall cell cycle shortening.

The cell cycle length analysis in the proliferative growth phase revealed that the rate of cell proliferation varied around an average value of ~0.11 cell divisions per cell per hour (Fig 2G, S3 Fig). Furthermore, our measurements showed that at 48 hpf ~18,000 cells inhabit the retina (Fig 1C). However, if all cells were continuously dividing at an average rate of 0.11/h starting from 2,177 cells at 20 hpf, the number of cells at 48 hpf should reach ~47,000. To understand this discrepancy in cell number at 48 hpf, we analyzed the fraction of cells that exited the cell cycle. We assessed the number of differentiating cells between 36 hpf (the stage when neurogenesis starts[22]) and 48 hpf (S2 Fig) using FACS-sorting of the SoFa transgenic line, that labels all emerging retinal neurons[23]. This analysis enabled us to account for the number of neurons and total number of cells with a model where progenitors divide at a constant rate of 0.11/h. In this model all divisions give rise to progenitors prior to 36 hpf while 35% of the divisions giving rise to committed precursors/neurons[22,24] after 36 hpf (Fig 2H and Supplementary Notes). Therefore, using both, cell division and differentiation rates calculated from our data, we were able to correctly predict the total number of cells at different developmental stages of the retinal PSE (Fig 2H).

We next used tissue volume, cell volume and cell division rate measurements to decompose the volume change in the tissue into (i) changes of average cell volume and (ii) changes in the number of cells through cell division. The cumulative rate of changes of these quantities are shown in Fig 2B. We verified that the sum of the three contributions add up to 0 within the experimental standard deviation, thus validating our measurements. We found that, because the rate of growth is about twice as low as the overall rate of cell division, the average cell volume is decreasing at the same rate as the tissue volume is increasing (Fig 2B).

Taken together, our data show that overall, the retinal PSE increases its volume homogeneously and isotropically in the plane of the epithelium, and cell division rates are about twice as large as the volume growth, leading to an exponential decrease in average cell volume.

### Retinal PSE growth is not limited by apical surface availability

The combined decrease in cell volume and increase in cell number lead to increased cell density in the retinal PSE (Fig 1B, S2 Fig). We wondered whether this increase in tissue density could influence growth dynamics in the retina through a self-inflicted proliferative trap. Such a proliferative trap was already proposed in the 1970s to arise from the fact that cell divisions occur exclusively at the apical surface [15,17,25]. This implied that when too many nuclei are packed under the apical surface or too many cells divide simultaneously, the availability of the apical surface becomes a spatial limitation to further proliferation. To test this idea, we used our quantitative dataset (Fig 1 and 2) and analyzed the fraction of the apical surface occupied by rounded mitotic cells between 20 hpf and 48 hpf. This analysis showed that mitotic cells never occupied more than 20% of the total apical surface, even at times of highest mitotic activity (S2 Fig). This result was confirmed by calculating the maximal number of proliferative cells in the volume beneath a single mitotic cell, approximated as a truncated cone to account for the wedge-shaped retinal cells (Fig 3A). Here, measured values did not exceed 40% of the calculated stacking maximum (Fig 3B), again suggesting that no proliferative trap exists in the retinal PSE.

**Figure 3.**
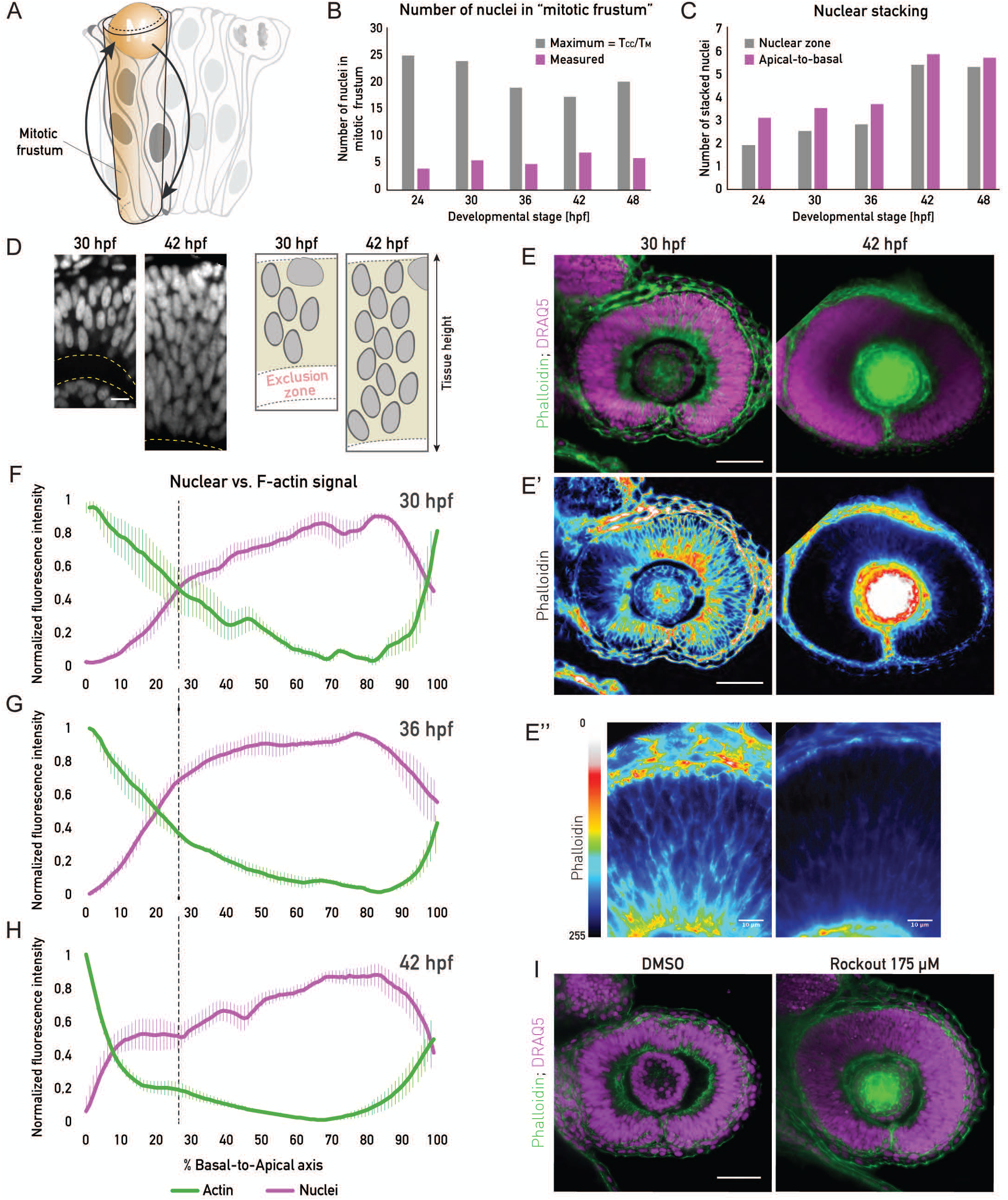
A tissue-wide basolateral actin accumulation establishes a basal nuclear exclusion zone. **(A)**, Schematic representation of PSE tissue architecture, with apical mitoses, migrating nuclei (arrows) and the mitotic frustum. The mitotic frustum is depicted as a conical unit below the rounded mitotic cell. **(B),** Number of cells under the rounded mitotic cell. Measured values are calculated from tissue-wide nuclear density and mitotic frustum volume in each developmental stage (10 samples/stage). Maximal possible number of cells is calculated as cell cycle length (T_CC_) divided by the duration of mitosis (T_M_) in each stage (see Methods). **(C)**, Number of stacked nuclei along the apico-basal axis of the retinal PSE. Number in the nuclear zone was calculated from nuclear stainings, dividing the thickness of the nuclear signal zone by the average long axis of the nuclei. Apical-to-basal number of layers was calculated as total tissue height/average nuclear long axis. **(D)**, Image and illustration of the basal nuclear exclusion zone in the first growth phase (30 hpf) and its absence in the second growth phase (42 hpf). Nuclear staining: DRAQ5. Scale bar: 10 µm. **(E)**, Tissue-wide basolateral actin accumulation (phalloidin) and the nuclear zone (DRAQ5) at 30 and 42 hpf. e’) and e’’) Lookup table indicates minimal and maximal phalloidin signal values. e’’) zoom-in on apico-basal tissue axis in e’). Dashed lines indicate apical and basal tissue surfaces. Scale bar in e and e’: 50 µm. Scale bar in e’’: 10 µm. (**F), (G), (H)**, Normalized average intensity distributions of phalloidin (darker shade) and DRAQ5 (lighter shade) signal in the tissue volume along the apico-basal axis of the retinal PSE at 30 hpf (**F**), 36 hpf (**G**), and 42 hpf (**H**). Dashed black lines indicate border of basal exclusion zone/ actin accumulation at 30 hpf. Values in each sample are normalized to minimum and maximum. Data is shown as mean ± SEM. 5-9 samples/stage. **i**, Rockout-treated and 0.3% DMSO-treated retinal PSE at 42 hpf. 175 µM Rockout treatment was started at 30 hpf.

Thus, proliferative retinal growth is not constrained by apical surface availability and a proliferative trap [17,19,25,26] does not apply to all growing pseudostratified tissues.

### A basolateral actin bias establishes a nuclear exclusion zone

While proliferative growth was not constrained by the apical surface, the number of nuclei stacked in the tissue nevertheless increased during the proliferative phase (Fig 3C). We analyzed this increase in nuclear stacking throughout development by dividing the height of the entire nuclear zone by the average nuclear height (Fig 3C, D). Between 36 hpf and 42 hpf nuclear stacking increase was most prominent, with the average number of stacked nuclei rising from 2.8 to 5.5 nuclei, respectively (Fig 3C, grey bars).

The tissue height measured in the nuclear channel, however, differed from the height in our initial growth analysis (Fig 1H) measured using the membrane channel (Fig 3C). One explanation for this could be that nuclei do not occupy the entire length of the tissue. Inspecting the tissue in more detail, we indeed noted that, prior to the significant increase in nuclear stacking, the basal zone of the retinal PSE was devoid of nuclei (Fig 3D). This resulted in a shorter nuclear zone compared to the entire tissue thickness (Fig 3D). This nuclear exclusion zone was reduced by 42 hpf, the time at which the major increase in nuclear stacking is observed. From this stage on, nuclei occupied the entire apicobasal tissue axis (Fig 3C, D).

We next asked how nuclei were restrained from occupying the basal region before 42 hpf. It was previously noted that a basolateral actin accumulation existed in the early optic cup [2]. Therefore, we tested whether this actin accumulation persisted at later stages and was linked to the basal nuclear exclusion zone. To this end, we analyzed the distribution of actin signal intensity along the apicobasal axis of the cells, in relation to nuclear distribution (Fig 3F-H). Before 40 hpf, the basolateral actin accumulation was consistently observed (Fig 3E, E’’, F). More specifically, at 30 hpf the basolateral actin accumulation took up 26% of the total cell height, or an average 12 µm of the apico-basal cell axis (Fig 3F). However, at 36 hpf the accumulation started diminishing (Fig 3G), spanning an average 20% of the cell axis (9 µm). At 42 hpf the basolateral actin pool was drastically reduced (Fig 3E, E’, E’’, H), occupying only the most basal 8% (5 µm) of the cells’ apico-basal axis. At all stages, this tissue-wide basolateral actin accumulation was in precise negative correlation with the nuclear exclusion zone (Fig 3F-H).

To test whether this actin accumulation was indeed responsible for the exclusion of nuclei from basal positions, we abolished basolateral actin at 30 hpf using the Rho-kinase inhibitor, Rockout [2]. This treatment led to a disappearance of both the actin accumulation and the basal nuclear exclusion zone, as nuclei were consequently seen along the entire apicobasal axis (Fig 3I).

Thus, basolateral actin accumulation creates a basal nuclear exclusion zone and only once it disappears nuclei occupy basal positions in the tissue.

### Actin redistribution is linked to an increase of cell height

The redistribution of actin was not only linked to an increase in nuclear stacking, but also coincided with the cell height increase observed after 36 hpf (Fig 1H, 4A). As contraction and tension of the actin cytoskeleton can actively affect cell shape in many different developmental contexts [27–29], we asked whether actin redistribution was involved in cell elongation. To test this notion, we analyzed actin redistribution in cells by comparing the ratios of the lateral signal intensity to basal epithelial belt actin signal intensity, throughout the proliferative phase (Fig 4A, B). We found that, as development progressed, actin was depleted from the lateral cell cortex and concentrated at the basal cell attachment (Fig 4B). If actin distribution was indeed involved in controlling tissue height, one simple explanation could be that the changing cell aspect ratio is dictated by a balance of forces between the line tensions in the apical and basal actin epithelial belt and the surface tension, exerted by the actin cortex along the lateral cell interfaces (S3 Fig and Supplementary Notes). The actin redistribution observed between 30 hpf and 42 hpf would then drive cell height elongation by favoring a contraction of the apical and basal perimeter at the cost of an expansion of the lateral surface area. We incorporated this possibility in our simplified description of tissue growth by considering a mechanical balance equation that gives the cell shape as a function of the lateral surface tension and the apico-basal line tensions (S3 Fig, Supplementary Notes). We also proposed an overall description of tissue growth with a constant rate of volume change and a constant rate of progenitor cell divisions. We further assumed that a transition occurs at ~35-36 hpf, at which progenitors start to differentiate, and actin redistributes from the lateral surfaces to apical and basal epithelial belts (Fig 4C, D). Taking into account the balance equations discussed earlier (Fig 2 and Supplemental Notes), this simplified description accounted for the main aspects of the geometrical changes seen for the tissue between 20 hpf and 48 hpf (Fig 4E-G).

**Figure 4.**
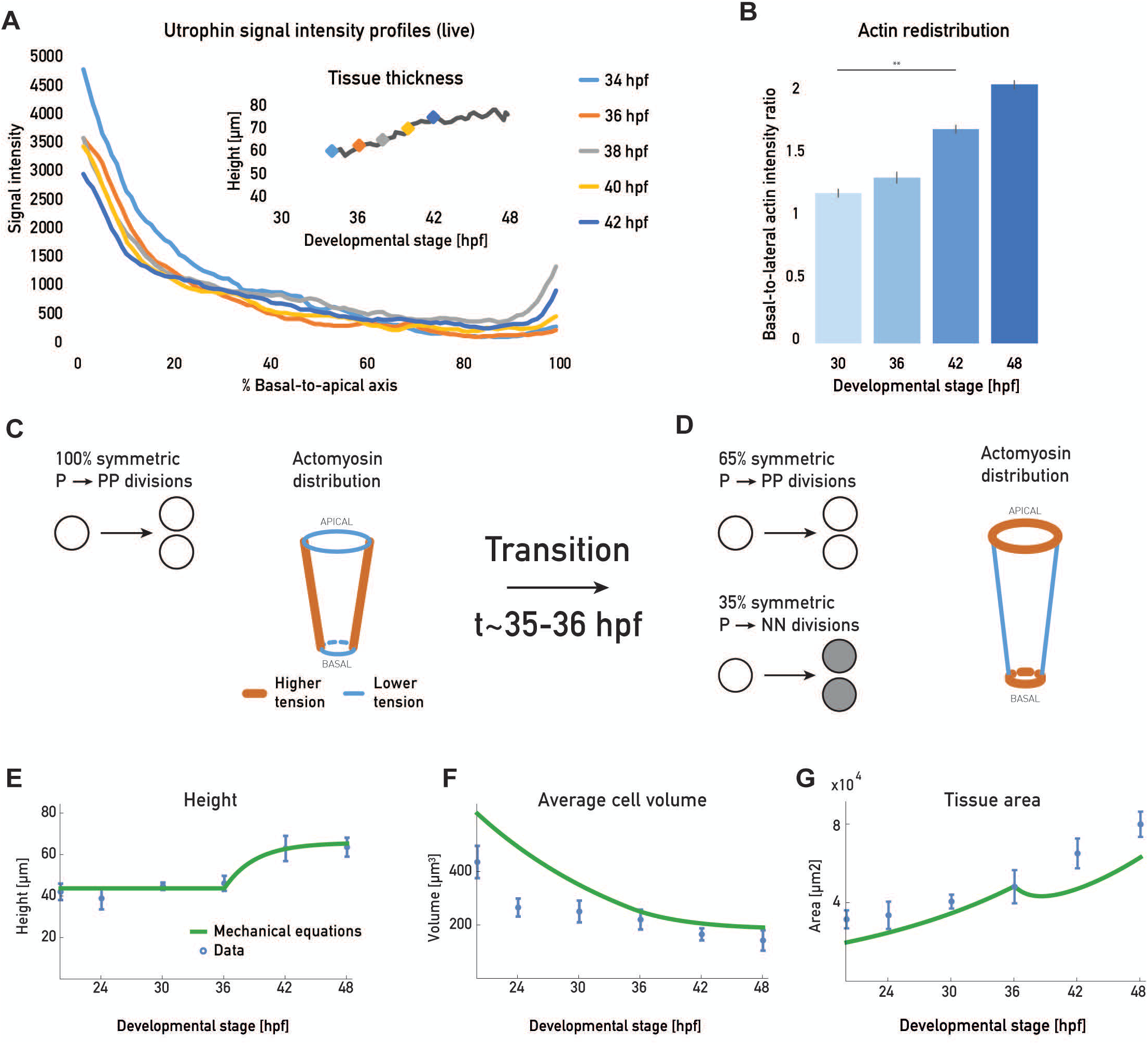
Basolateral actin accumulation coincides with cell height increase. **(A)**, Apico-basal actin intensity distributions from live Tg(actb1:GFP-UtrophinCH) embryo measurements over development. Inset plot shows cell height increase in the same sample, with actin intensity measurement points marked (color code matches main plot) (S6 Movie). **(B)**, Actin redistribution over development. Ratios of basal-to-lateral phalloidin signal. 5 samples/ stage. Mean ± SD. Mann-Whitney test, p-value 0.0079. (**C), (D)**: Schematic of simplified theoretical description of the retina growth. Prior to ~35-36 hpf, all progenitor divisions give rise to two progenitors, and a constant ratio of apicobasal line tension to lateral surface tensions is maintained. After this transition, 35% of the divisions give rise to committed progenitors or neurons, and a redistribution of actin drives a change in the ratio of apicobasal line tension to lateral surface tension. **(E), (F),(G)**: Blue dots: experimental data, plotted as mean±SD. green line: theory prediction for height, cell volume and tissue area.

To experimentally test whether a link between actin redistribution, cell height increase and uniform tissue growth existed, we used a histone-deacetylase 1 (hdac1) mutant, shown to undergo continued proliferation and not enter differentiation programs due to the loss of Hdac1-dependent Wnt and Notch inhibition [30,31]. Continued proliferation ensues without isotropic shape scaling and the hdac1-/- retinal PSE exhibits a perturbed shape[30,31] (S4 Fig). Live imaging confirmed that hdac1-/- mutants developed a perturbed retinal PSE instead of the smooth tissue surface seen in controls (S4 Fig, S4 Movie). This shape disruption starts at stages at which differentiation would have begun in the wild-type retina (S4 Fig, S4 Movie). In previous studies, the emergence of this shape distortion was not explicitly explored, but it was reported that hdac1-/- mutants show disturbed actin organization[31]. We thus set out to test whether a lack of actin redistribution could be linked to the shape disruption seen in this mutant. We imaged actin localization and cell height changes of control versus hdac1-/- neuroepithelia (Fig 5 and S5 Movie). This revealed that in hdac1-/- embryos, actin accumulation along lateral interfaces lasted substantially longer than in controls (70 hpf vs. 40 hpf, Fig 5A-C). This was confirmed by the analysis of basal-to-lateral actin signal ratios, which, unlike the control, in the hdac1-/- remained unchanged throughout development (Fig 5D).

**Figure 5.**
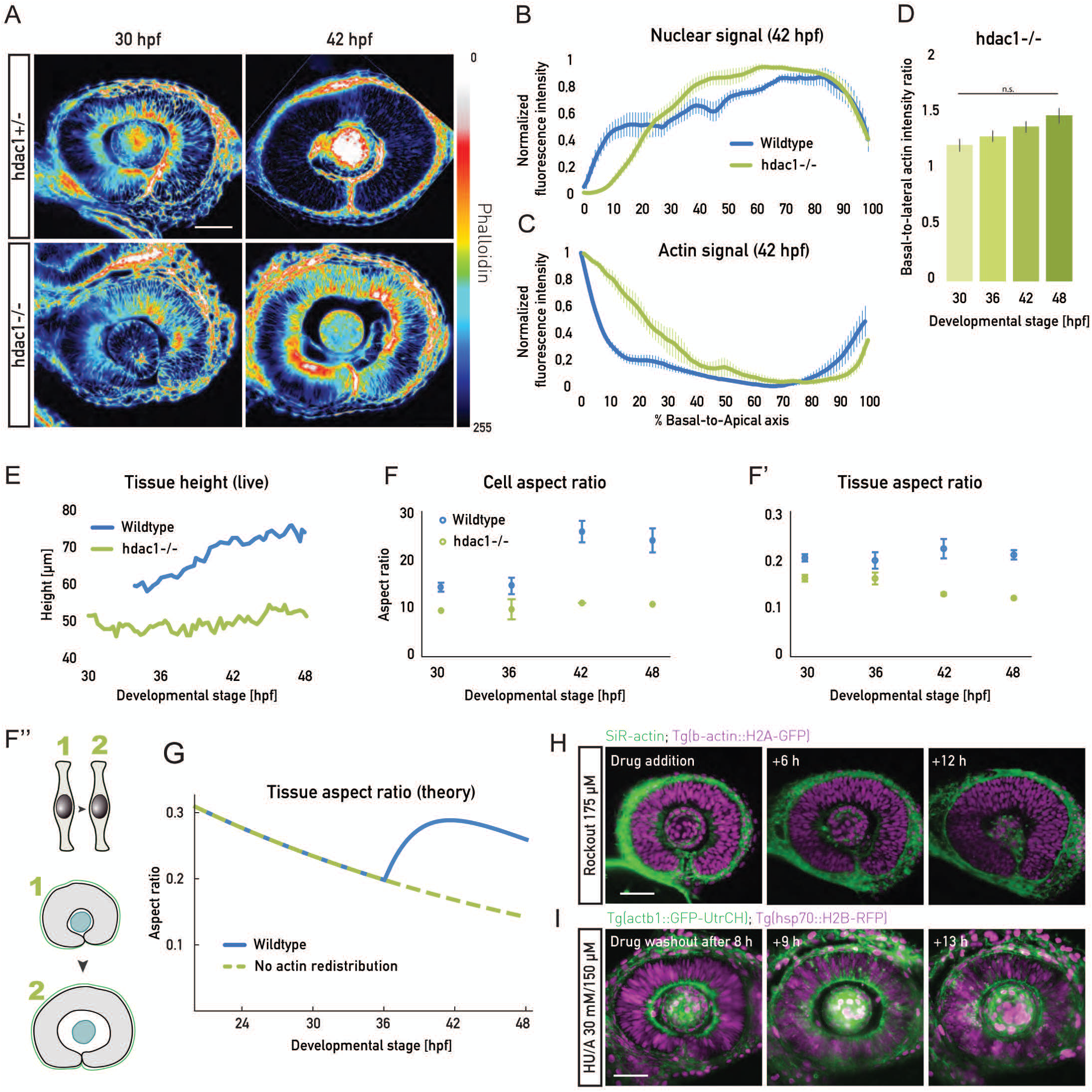
Actin redistribution allows for cell elongation in concert with cell proliferation. **(A)**, Phalloidin signal before (30 hpf) and after (42 hpf) actin redistribution in the wildtype and heterozygous hdac1+/- controls. Basolateral actin accumulation does not redistribute in hdac1-/- samples (bottom panels). Lookup table indicates minimal and maximal phalloidin signal values. **(B), (C),** Normalized average intensity distributions of DRAQ5 (**B**) and phalloidin (**C**) signal in hdac1-/- (green) and control (blue) samples at 42 hpf. Values in each sample are normalized to minimum and maximum values. Data is shown as mean ± SEM. 3-9 samples/stage. **(D)**, Basal-to-lateral phalloidin signal intensity ratios in hdac1-/- samples. 5 samples/stage. Mean ± SD. Mann-Whitney test, p-value 0.3095. **e**, Tissue height measurements from live embryos (light sheet time-lapses) for control (blue) and hdac1-/- (green) samples (see S5 Movie). **(F)**, Cell shape analyzed as aspect ratios from mean cell cross-sectional area and height in control (blue) and hdac1-/- (green) samples. 2-10 samples/stage. **(F’)**, Tissue shape analyzed as aspect ratios from mean tissue area and height in control (blue) and hdac1-/- (green) samples. 2-10 samples/stage. (**F’’)**, Schematic representation of the unchanged cell aspect ratio during hdac1-/- retinal PSE growth and perturbed tissue shape (see also S4 and S5 Movies). **(G)**, Prediction of simplified theory for the tissue aspect ratio 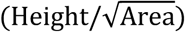 in the model described in Fig 4C-D (blue line) and in a model where the actomyosin does not redistribute from apicobasal junctions to the lateral cortex (green dotted line, see Supplementary Notes for details, compare with Fig 5F’). **(H),** 175 µm Rockout treatment abolishes the basolateral actin accumulation but is insufficient to induce cell elongation. Scale bar: 50 µm. Actin is labelled by SiR-actin (green), and nuclei by Tg(b-actin:H2A-GFP). **(I)**, Proliferation is necessary for cell elongation. After HU/A treatment (30 mM HU + 150 µm A) cell proliferation is blocked, but cell height does not increase. Actin is labelled by Tg(actb1:GFP-UtrCH) (green), and nuclei by Tg(b-actin:H2B-RFP) (magenta).

When the basolateral actin bias was maintained in hdac1-/- cells, cells did not elongate past 50 µm (Fig 5F), a height value that is reached in controls at 30 hpf. As a consequence of hdac1-/- cells not changing their shape (Fig 5F, F’), the retinal PSE did not maintain its aspect ratio upon continued proliferation (Fig 5F’, F’’, S5 Movie) leading to tissue shape changes instead of cell shape changes. This indeed indicated a link between retinal PSE shape changes and actin rearrangements. In accordance with this finding also our simplified theory predicted that the lack of actin redistribution leads to an excess decrease in tissue aspect ratio as seen in the hdac1-/- mutant, (Fig 5G).

While PSE shape was disturbed in the hdac1-/- neuroepithelium, the tissue nevertheless retained its pseudostratified characteristics, as shown by intact apical junctions and basal lamina (S4 Fig), and its mitoses occurring apically (S4 Fig). A similar sequence of tissue shape changes was seen when using an Hdac1 morpholino (S4 Fig) [30] or the Hdac1 inhibitor Trichostatin-A (S4 Fig). In all conditions, the end result of shape perturbations was the appearance of apical and basal epithelial folds (S4 Fig) [30,31].

Taken together, this data suggests that actin redistribution from lateral cellular interfaces to the apical and basal epithelial belts drives cell and thereby tissue elongation. When this reorganization does not occur, cells do not increase their height, leading to a disturbed tissue shape.

### Actin redistribution allows for tissue height increase in concert with cell proliferation

We asked whether the redistribution of actin from the lateral cellular interfaces to the epithelial belts is sufficient to drive cell elongation. To test this, we used Rockout to deplete the lateral actin accumulation prematurely (Fig 5H). As a result, nuclei redistributed along the apico-basal axis and filled basal positions (Fig 5H). However, even after the basal actin accumulation was diminished and nuclei inhabited basal positions, no instantaneous increase in tissue height was observed (Fig 5H). This argued that actin disappearance from lateral cellular interfaces was necessary, but not sufficient to trigger cell elongation, suggesting that additional factors contribute to PSE thickening.

As the increase in cell height observed after 36 hpf coincided with increased proliferation due to shortened cell cycles (Fig 2 F, G), we tested whether proliferation is involved in cell and tissue elongation. To this end, we blocked proliferation using HU/Aphidicolin treatment at 30 hpf. In this condition, even after the lateral actin accumulation disappeared, the tissue height did not increase to the same amount as in controls (Fig 5I, S5 Fig).

Together, these experiments suggest that both the disappearance of actin from lateral interfaces and continued proliferation are required for the observed cell and tissue elongation. Thus, these two factors together ensure isotropic scaling during retinal PSE growth.

### Actin distribution reorganizes non-cell autonomously and is dependent on the extracellular matrix

Live imaging of F-actin revealed that the disappearance of actin from lateral interfaces occurred simultaneously throughout the retinal PSE (S6 Movie). This indicated that actin rearrangements occur in a non-cell autonomous manner. To test this idea, we transplanted cells from control embryos into hdac1-/- embryos, in which the basolateral actin bias persisted (Fig 5A and C). Here, transplanted control cells preserved their actin distribution and did not increase their height, even at stages at which they would elongate in their native environment (Fig 6A, A’). Conversely, when hdac1 morpholino was injected into 64-cell stage control embryos, to achieve a mosaic distribution of morphant cells, these morphant cells lost the lateral actin accumulation similarly to neighboring control cells (Fig 6B). In this context, nuclei of hdac1 morphant cells occupied basal positions and cells increased their height alongside their neighbors (Fig 6B). This confirmed that actin redistribution is a non-cell autonomous event, linked to cell height increase.

**Figure 6.**
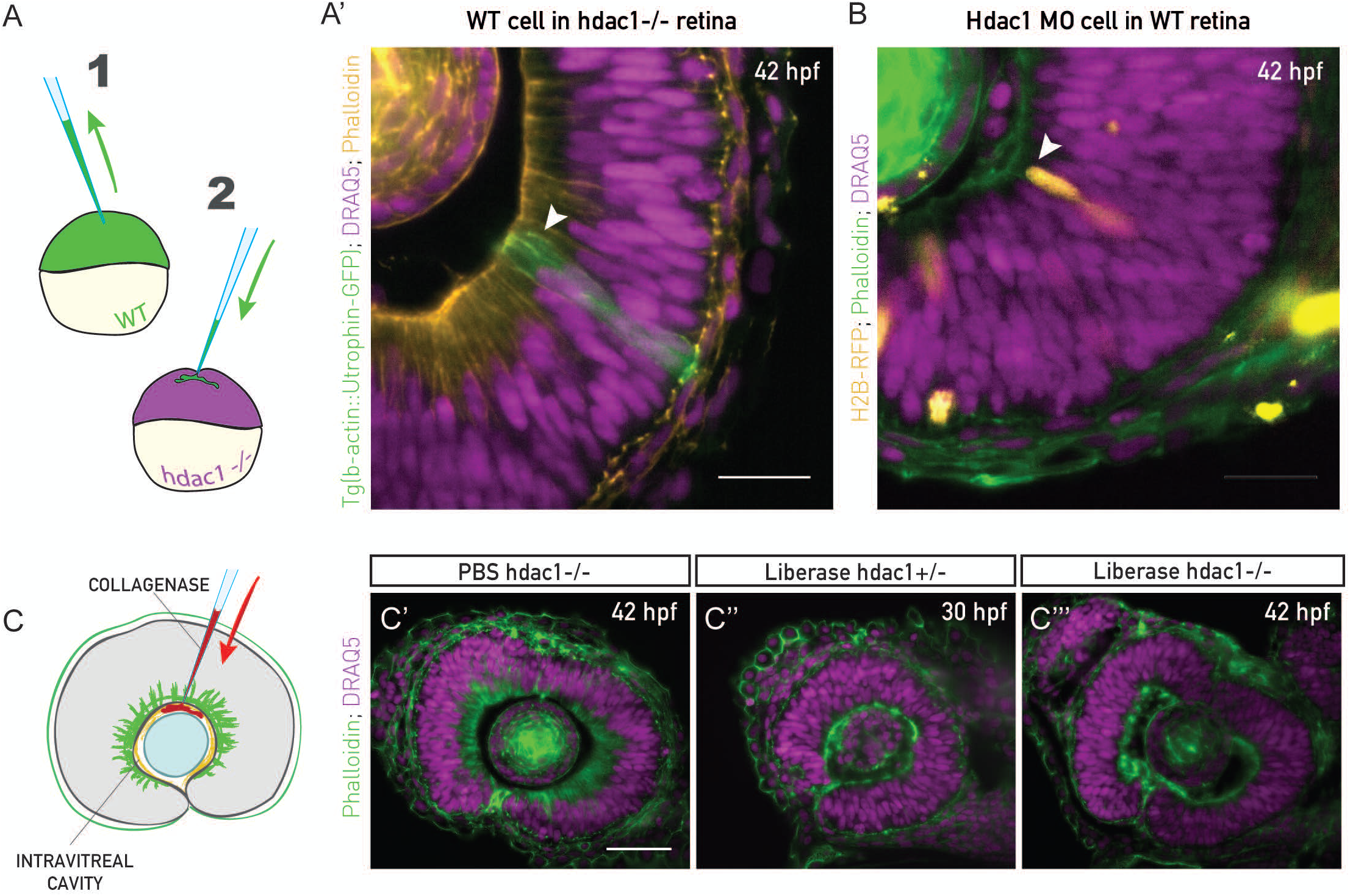
Actin reorganization and cell height increase are non-cell autonomous and ECM-dependent. **(A)**, Schematic of the transplantation experiment. Blastomeres from Tg(actb1:GFP-UtrCH) embryos (1) were transplanted into acceptor hdac1-/- embryos (2) between high and sphere stage. **(B)**, WT cells in hdac1-/- tissues do not redistribute lateral actin nor increase their height at 42 hpf (green cells, arrowhead). Actin is labelled with Phalloidin (orange) and nuclei with DRQ5 (magenta). N = 5 embryos. **(B’)** Mosaic injection of hdac1 morpholino (MO) into Tg(actb1:GFP-UtrCH). Cells with hdac1 MO were co-injected with H2B-RFP mRNA for visualization. Their basal actin zone disappears at the same time as in the WT (orange cells, arrowhead). Actin is labelled with Phalloidin (green) and nuclei with DRAQ5 (magenta). N = 5 embryos. **(C)**, ECM integrity was perturbed by injecting 0.5 mg/ml of collagenase (Liberase) into the intravitreal cavity. **(C’)**, The basal actin accumulation in hdac1-/- tissues remains intact when PBS is injected as control. **(C’’)**, Collagenase abolishes the basal accumulation in 30 hpf control hdac1+/- embryos. **(C’’’)**, Collagenase abolishes the basal actin accumulation in 42 hpf hdac1-/- tissues. C’-C’’’) actin is labelled with Phalloidin (green) and nuclei with DRAQ5 (magenta).

It has been previously shown that the ECM underlying developing tissues can actively influence tissue size and shape during organ development [32,33]. The retinal PSE cells are basally attached to the ECM of the basal lamina, that contains collagen and laminin as main components (S4 Fig) [34]. As forces mediated by cell-ECM attachments can affect actin polymerization and bundling in diverse epithelial cells [35,36], we tested whether cell-ECM interactions were involved in basal actin reorganization in the retinal PSE. We injected collagenase, an ECM-degrading metalloprotease, between the lens and the retinal neuroepithelium of hdac1+/- and hdac1-/- fish at 30 hpf and 42 hpf (Fig 6C). In the hdac1+/- control, as well as in the hdac1-/- tissues, collagenase indeed abolished the actin accumulation on lateral interfaces and led to the removal of the nuclear exclusion zone (Fig 6C’’, C’’’) strongly suggesting an active role of the ECM in actin reorganization. Thus, ECM composition influences isotropic growth of the retinal PSE.

This data shows that actin redistribution is indeed a prerequisite for cell height increase, that occurs independently of cellular identity. In addition, actin redistribution is dependent on an intact ECM.

## Discussion

In this study, we decomposed growth of the retinal PSE in geometric cell and tissue-wide parameters and show that proliferative retinal growth is uniform and isotropic. We find that this uniform 3D growth is enabled by timely redistribution of the actin cytoskeleton in concert with continued proliferation, allowing for timely apicobasal cell elongation.

To understand the links between growth and shape in a developing PSE in 3D, we generated a quantitative dataset of the retinal neuroepithelium. This, together with a set of balance equations, allowed us to generate an unprecedented characterization of growth. We reveal that geometric changes at the single cell level are essential to maintain tissue shape. Single cell shape changes were most prominent after 36 hpf, a stage at which tissue height increased.

In concurrence with this increase in tissue height, nuclei enter a previously unoccupied nuclear exclusion zone, leading to an increase in nuclear stacking. We find that the loss of this nuclear exclusion zone, as well as cell height increase, are linked to a cellular redistribution of actin. When actin is maintained at lateral cell interfaces, cell elongation does not take place and the isotropic scaling of tissue shape is consequently disturbed. These findings let us speculate that the lateral actin accumulation generates a mechanical force acting against apico-basal cell elongation. We postulate that the redistribution of actin leads to a change of forces generated in the cells that are involved in driving apicobasal cell elongation. How exactly height increase results from the observed actin redistribution will need to be investigated in future studies.

Basolateral actin accumulation depends on an intact underlying ECM, as we found that disturbing ECM composition results in actin redistribution. This together with the fact that actin redistribution occurs non-cell autonomously supports the idea that ECM rearrangements or cell-ECM attachments act as upstream modulators of actin cellular organization. Such changes could be driven for example by the Wnt or Notch pathway, as Hdac1 acts as inhibitor of Wnt and Notch and both molecules have been implicated in modulating cell-ECM force transmission [37–39]. The thorough analysis presented here will be an important baseline for future studies to investigate these factors and their involvement in cell shape changes and tissue shape maintenance.

Our results also demonstrate that tissue growth needs to be tightly coordinated to cell shape changes in time and space for successful tissue development. We show that cell height increase occurs in concert with the neighboring tissue independently of how the cell would behave in its native environment. In contrast, retinal neuronal differentiation is most likely a cell autonomous program as we observed that after washing-out the hdac1 inhibitor Trichostatin-A, cells in the already folded retinal tissue regained the potential to differentiate and generate neuronal layers, in spite of the tissue’s perturbed shape (S4 Fig). Thus, it is likely that differentiation is uncoupled from tissue growth. This lends further weight to the fact that the precise coordination of cell intrinsic and tissue wide parameters is essential to generate the final, differentiated tissue of correct size and shape.

We found that in the retinal PSE, apical surface availability does not pose a constraint on growth during the proliferative phase, as was suggested for other PSE [15,17,25]. While it is possible that the retinal PSE is an exception due to its curved shape, a careful analysis of factors contributing to tissue growth in other PSE, as done here, would allow to test this old hypothesis [17] in other contexts.

As retinal cells need to change their geometry in concert with proliferation, our study exemplifies that tissue shape maintenance during growth is not a default state but requires active preservation [40]. We found that isotropic shape scaling of the retina depends on altering active forces in the cell which control the cell shape. Such dynamic force change is also seen in other contexts such as 2D germ band elongation in Drosophila [4]. In our example, however, this force redistribution acts in 3D along the apicobasal axis of cells and ensures isotropic scaling of shape during growth, rather than changes in tissue shape.

Overall, we elaborate on the important question of how organs coordinate growth with isotropic scaling of shape during development. We believe that our measurements can constitute a reference point when assessing remaining questions of growth in diverse developing PSE. It has recently become more feasible to test such questions in developing embryos *in vivo* due to new and improving imaging techniques [41]. Quantitative studies as presented in here will need to be extended in the future using more model tissues and organisms. The next challenge will then be to expand investigations to growth phenomena of *ex vivo* tissues and organoid systems including human organoids, which often emerge from PSE, as seen for retinal and cerebral organoids [42–44]. In-depth 3D studies of growth and shape will then allow to compare growth phenomena of *ex vivo* and organoid cultures to *in vivo* growth. Such comparisons will deepen our understanding of developmental programs in model organisms and humans.

## Materials and methods

### Zebrafish husbandry

Zebrafish were maintained and bred at 26.5°C. Zebrafish embryos were raised at 21°C, 28.5°C or 32°C and staged in hours post fertilization (hpf) according to [45]. E3 medium was exchanged daily. 0.003% 1-phenyl-2-thiourea (PTU) was added to the embryo E3 medium from 8±1 hpf onwards, and renewed daily to prevent pigmentation. Embryos were anaesthetized in 0.04% tricane methanesulfonate (MS-222; Sigma-Aldrich) prior to sorting for fluorescent signal (>36 hpf), imaging, intravitreal cavity injections, brain ventricle injection or retinal dissection. All animal work was performed in accordance with the European Union (EU) directive 2010/63/EU as well as the German Animal Welfare act.

### Transgenic and mutant lines

See S1 Table for a list of genetically modified lines used in this study.

### Hdac1 mutant line

Hdac1 t24411 mutants, containing a single point mutation in the histone deacetylase 1 (hdac1) gene were obtained from the Zebrafish International Resource Center (ZIRC). Incrossed, homozygous fish (hdac1-/-) until 3 dpf were used in experiments.

### Blastomere transplantation

Donor embryos (Tg(actb1:GFP-UtrCH)) and acceptor embryos at stages high to sphere were dechorionated in pronase and cells from the animal pole were transplanted into the acceptor hdac1+/-, -/- embryos. Transplanted embryos were kept on agarose for ~3-5 h and then transferred onto glass dishes. The E3 medium was supplemented with antibiotics (100 U of penicillin and streptomycin, Thermo Fisher Scientific). Homozygous mutants were identified, fixed and stained after 30 hpf.

### Morpholino injections

Morpholinos were produced by Gene Tools, LLC.

Hdac1 MO was used to knock down Hdac1 mosaically, or in all cells. p53 MO was added to the morpholino injection mix to alleviate negative effects associated with injection and knockdown. All morpholino mixes were injected into the yolk of 1- to 2-cell stage embryos. See S2 Table for list of used morpholinos.

## mRNA and plasmid injections

To label all cells, mRNA was injected into the yolk or cell of 1-cell stage embryos (100 pg/embryo). The injection volume was 1-2 nl. mRNA was synthesized using the Ambion mMessage mMachine kit. To label cells mosaically, mRNA was injected into single cells of 32-128-cell stage embryos (50 pg/cell in 0.5 nl injection volume). Plasmid DNA was injected into the cell of 1-cell stage embryos (5-15 pg/embryo in 1-2 nl).

## Intravitreal cavity collagenase injections

To perturb the ECM locally, collagenase (Liberase, Sigma Aldrich), an ECM degrading metalloprotease, was injected into the intravitreal cavity (between the lens and the retina). 0.5 mg/ml Liberase in a volume to fill the cavity (2-3 nl) was injected in 36 hpf or 42 hpf hdac1-/- embryos, and 30 hpf control embryos. Injected fish were incubated in a waterbath at 37°C to activate the enzyme for 30’. Afterwards they were fixed and stained with phalloidin.

## Heatshock of embryos

Embryos injected with plasmids containing Hsp70:H2B-RFP, or transgenic embryos (Tg(Hsp70:H2B-RFP) were heat shocked at least 1.5 hours prior to imaging. Embryos injected with Hsp70:H2B-RFP plasmid were heat-shocked for 10’ at 37°C. Tg(Hsp70:H2B-RFP) embryos were heat-shocked for 20’ at 38°C.

## Drug treatments

The minimal effective concentrations of inhibitors were optimized based on dilution series. All inhibitors were dissolved in DMSO, except for hydroxyurea, which was dissolved in water. Dechorionated embryos were treated by incubation in the inhibitor-E3 medium at specific concentrations, either in plastic multi-well plates or directly in a light sheet microscope chamber (Rockout, Trichostatin-A). All treatments started after 24 hpf (Rockout in WT always before 42 hpf) and lasted from 3 h to 2 days (Trichostatin-A), with most treatments overnight (~15 h). See S3 Table for the list of used chemical inhibitors and concentrations.

## Immunostaining

Wholemount immunostainings were performed on pronase-dechorionated embryos, fixed in 4% paraformaldehyde (Sigma) in PBS, at +4°C overnight. Washes were performed in PBS-T (PBS with 0.2% or 0.8% Triton X-100). Embryos were permeabilized in 1% Trypsin, blocked in 10% normal goat serum (NGS), or blocking buffer [46]. Embryos were incubated in primary antibodies for 64 h, shaking at +4°C.

### Primary antibodies

The phosphorylated histone H3 (PH3) antibody (Abcam; ab10543, RRID:AB 2295065) was used at 1:500 to label chromatin of mitotic cells. Laminin α1 antibody (Sigma-Aldrich; L9393, RRID:AB_477163) and Collagen IV antibody (Abcam; ab6586, RRID:AB_305584) were both used at 1:100 to label the basal lamina.

### Secondary antibodies

Secondary antibodies with conjugated fluorophores Alexa Fluor 405, Alexa Fluor 488, Alexa Fluor 568 and Alexa Fluor 647 (Thermo Fisher Scientific Inc.) were used at 1:500 or 1:1000.

### Fluorophore-conjugated compounds

To label all cell nuclei, DAPI (1:5000, or 1:2500) and DRAQ5 (1:2500, Thermo Fisher Scientific Inc.; 62251) were used. Phalloidin-TRITC (1:50, Life Technologies; R415), Phalloidin-Alexa Fluor 488 (1:50, Life Technologies; A12379) were used to label F-actin.

## Cell dissociation and flow cytometry

To obtain relative numbers of differentiating cells in proportion to all retinal PSE cells, Tg(SoFa) retinas [23] were dissected, dissociated and analyzed by FACS.

### Retinal dissection and dissociation

Dissection was performed on Sylgard-coated 10 cm plastic dishes, at room temperature in PBS. Eyes were dissected from live 36 hpf, 42 hpf and 48 hpf Tg(SoFa) and 48 hpf wildtype anaesthetized embryos using forceps, syringe needles and/or a 5 mm surgical stab knife. The pigment epithelium and lens were removed. In total, 20 retinas/stage were transferred to a glass FACS-tube. Cells were dissociated mechanically and immediately analyzed by FACS.

### FACS-sorting

Single cell suspensions were analyzed using the FACSAria device. Wildtype cells were used to normalize the signal to background auto-fluorescence. Populations were defined based on forward- and side-scatter signals in PE-A, FITC and DAPI channels. Data was plotted and processed using the FACSDiva software. The resulting scatterplots were analyzed as described previously for the SoFa line[23], in order to obtain relative values for different neuronal subtypes. Total numbers of cells in the retina at different developmental stages were used to calculate average absolute numbers of differentiated cells from the relative values.

## Image acquisition

### Confocal laser point-scanning microscopy

Dechorionated, fixed and immunostained embryos were mounted in 1% low melting-point agarose in glass-bottom dishes (35 mm, 14 mm microwell, MatTek). The agarose dome was covered with PBS or E3. Imaging was performed on Zeiss LSM 510 or 710 confocal microscopes (Carl Zeiss Microscopy) using a 40x/1.2 NA water immersion objective, at room temperature. The imaging system was operated by the ZEN 2011 (black) software.

### Light sheet microscopy

Dechorionated live or fixed embryos were mounted in glass capillaries in 0.6% or 1% low-melting-point agarose. Imaging was performed on a Zeiss Light sheet Z.1 microscope (Carl Zeiss Microscopy) with a Zeiss Plan-Apochromat 20x or 40x water-dipping objective (NA 1.0) at 28.5°C. For drug treatment experiments, the chamber was filled with the drug-supplemented medium. For fixed sample imaging, embryos were mounted in 1% or 0.6% agarose, immersed in PBS and imaged at room temperature. Z-stacks spanning the entire eye were recorded with 0.5-1 μm optical sectioning. For live imaging, z-stacks were recorded every 5-30’. The system was operated by the ZEN black software.

## Image analysis

Minimal image preprocessing was implemented prior to image analysis. Processing consisted of extracting image subsets, bleach correction and/or background subtraction using Fiji[47]. XY-drift was corrected using a Fiji plugin (Manual Registration, Scientific computing facility, MPI-CBG). After image analysis in IMARIS 8 (BitPlane) or Fiji, data was analyzed and plotted using Matlab or Microsoft Excel. Statistical analysis was performed using the Prism software (version 6.0c for Mac OS; GraphPad Software).

## Growth analysis

Most image analysis was done on the entire tissue in 3D, using IMARIS 8. This included tissue and cell volumes, total tissue area, tissue height, cell and mitotic cell position and count analyses. Raw data was segmented by manually outlining the retinal and lens tissues. 30 to 50 z-sections were outlined in each sample to span the retinal tissue, and 10 samples/stage, in 6 stages were analyzed as part of the WT growth characterization. The segmented rendered surface was used to mask the raw data and create a subset on which further measurements were performed.

## Mitotic density distribution, Script 1

To assess the distribution and the density of mitoses at the apical surface in different stages, mitotic cell positions were detected and projected onto a 2D density heatmap, using a custom Matlab tool, “PSE MitoNuc” (Benoit Lombardot, Scientific computing facility, MPI-CBG). pH3+ mitotic cells in the masked retinal image were detected in IMARIS using the Spot detection tool. Cartesian coordinates of all detected spots were exported from IMARIS and analyzed by the “PSE MitoNuc” tool. Briefly, the 3D mitotic coordinates were first transformed into spherical coordinates, and then projected onto a 2D surface using an existing density-preserving, azimuthal projection tool. To use the “PSE MitoNuc” tool, refer to the readme file in the supplement (README_Script1.txt). This method was validated on terminal neurogenic divisions (Ath5+), and the typical non-uniform distribution of divisions (naso-temporal wave) was clearly detected here (S2 Fig).

## Division orientation

Division orientations were analyzed manually in Fiji from live images of all cells labelled with Tg(actb1:HRAS-EGFP) (membrane). A part of the curved apical surface was imaged using the light sheet microscope. Data was bleach-corrected and background was subtracted. For each observed anaphase cell, a line was drawn perpendicular to the division plane and the angle was measured. Data was processed (to be between 0° and 180°) and plotted as a rose plot in MATLAB.

## Apical mitotic occupancy

Average cross-sectional areas of mitotic cells were calculated as circle areas from diameters of mitotic cells measured in fixed samples (20-40 cells/stage, 3-7 embryos/stage). This number was multiplied by the number of mitotic cells in each stage to obtain the total area occupied by mitotic cells. This area was divided by the total apical surface area of the tissue to obtain the fraction of the apical surface occupied by mitotic cells in each stage.

## Nuclear packing

For each developmental stage, the volume of the conical unit under the mitotic cell was calculated as the frustum volume

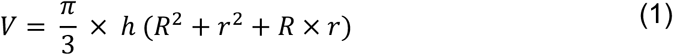

where h is the mean cell (tissue) height, R the mean radius of a mitotic cell and r the mean radius of the basal cell area. Mitotic cells cross-section and basal cell attachment are assumed to be circular. The cell volume density (cells/µm^3^) in the tissue was calculated by dividing the total number of cells by the tissue volume. The number of nuclei (cells) within the mitotic frustum was calculated by multiplying the cell volume density by the volume of the frustum.

Due to the apical localization of mitoses in the PSE, the average cell cycle time T_CC_ (time for which a nucleus is not rounded at the apical surface), needs to be longer or equal to the duration of mitosis T_M_ (time for which a cell is occupying the apical surface) of all cells in the frustum unit, N [17]

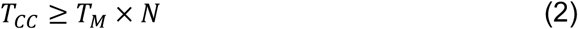

From Eq. 2, the maximal number of cells in the frustum unit depends on the cell cycle parameters and is equal to

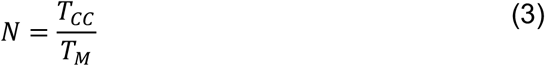

## Specific growth and shape parameters

### Apical and basal areas

Apical tissue area of the retinal PSE was measured using a custom Fiji tool (Volume Manager, by Robert Haase, Scientific computing facility, MPI-CBG). Areas were manually outlined. Analysis and visualization were performed using Volume Manager. Volume Manager is available from the Fiji SCF-MPI-CBG update site. Basal tissue area was calculated by subtracting the apical tissue area from the total tissue area.

Average apical and basal cell areas were calculated by dividing the apical and basal tissue area by the total number of cells, respectively.

### Cell size measurements and validation

Cell size was estimated using rounded mitotic cells. Diameters of 15-20 mitotic cells/retina (5-10 samples/stage) were measured using the IMARIS measuring tool. Assuming that mitotic cells are nearly spherical, cell volumes were calculated as sphere volumes. To check how well volumes of mitotic cells represent interphase cells, several interphase cell volumes were manually segmented from images with extremely sparse mosaic labelling of cells with GFP-Ras mRNA (cell membrane marker). This analysis indicated that mitotic cell volumes are on average 10% larger than interphase cell volumes, and the mitotic cell volume measurements were adjusted accordingly.

### Cell number validation

Total cell numbers were analyzed using the IMARIS spot detection tool to detect individual nuclei on the masked retinal stacks, from the DRAQ5/DAPI channel. The automatic detection was validated manually, by examining individual z-planes using the Oblique slicer tool and a combination of two Clipping plane tools to isolate and examine a specific tissue region. Automatic settings (thresholds) were manually adjusted for each sample. To validate the count, cell volumes were estimated from mitotic cell volumes. The total tissue volume was divided by these cell volumes, in order to obtain the average volume occupied by a single cell (i.e., cell volume). Cell volumes obtained in this way were in very good agreement to the values obtained by the automatic count (S1 Fig).

### Tissue thickness

Fiji was used to manually measure tissue thickness in live images of WT and hdac1-/- fish. The apicobasal tissue axis at the nasal region of the retina was measured at every time-point using the line tool. For fixed samples, thickness was measured in the central slices in the nasal and temporal region and proximal region (behind the lens). To correct for the lower thickness in the ciliary marginal zone, all tissue thickness values were decreased by 5%.

### Nuclear stacking

The number of nuclei stacked in layers in the nuclear zone was calculated by dividing the average height of the nuclear zone of the tissue measured from nuclear stainings, by the average length of the long (apico-basal) axis of the cell nuclei (30 nuclei/stage, 2 embryos/stage). The number of stacked nuclei along the entire apico-basal tissue axis was calculated at every stage by dividing the tissue height by the long axis of the nuclei. In addition to this, the number of layers was counted manually. These values were in very good agreement to the calculated values for the nuclear zone.

### Actin and nuclear signal intensity distribution, Script 2[2]

The average intensity distribution of phalloidin and DRAQ5 signal along the apico-basal axis of the PSE cells was measured as described in[2], using a custom Python script (Benoit Lombardot and Robert Haase). The region of interest was defined as a 10 μm x 10 μm x thickness cuboid (3-9 samples/stage). An average intensity value for this region was calculated for each point along the apico-basal axis. The axis length was normalized to 100. To compare different samples, the average intensities were normalized to the minimal and the maximal intensity value along the axis.

### Actin signal intensity ratios

To obtain the ratios of acting signal intensity, the apical, basal and lateral portions of the tissue were manually outlined using the Fiji line tool, in the central section of the retina for 5 samples/ stage. Apical and basal surfaces were outlined with 10 pt thick lines. Lateral regions were selected using 200-350 pt thick lines in the nasal, central and temporal part separately. Intensity profiles were processed to extract peaks of signal, assumed to represent the cortical signal, using the MATLAB *findpeaks* function.

### Cell cycle analyses

Mosaically labelled cells in long (>15 h) time lapse light sheet datasets, with time resolution of 5 min were manually tracked in 3D, in ZEN and Fiji. Cells were labeled with Hsp70:H2B-RFP. For total cell cycle analysis, time of chromosome segregation for the first, second, and, in wherever possible, third division was recorded. For the analysis of mitosis length, endpoints of mitosis were recorded as the point of cell rounding and the point of chromosome segregation. This cell cycle data for 254 cells from 20 embryos was analyzed, binned into discreet stages and plotted in MATLAB.

## Supporting information

Supporting Tables 1-3

Supporting Figures 1-5

Supplementary Notes

## Author contribution

Conceptualization, M.M., G.S., C.N., Methodology M.M., G.S., C.N., Investigation, M.M., G.S., Writing, C.N., M.M., G.S. Visualization, M.M., Supervision, C.N., G.S.

## Acknowledgements

We thank the Norden lab, W.A. Harris, C. Modes, J. Tabler and J. Vermot for helpful comments on the manuscript. S. Grill, A. Hyman, B. Langer, C. Modes, J. Tabler and the whole Norden lab are thanked for useful project discussions. We are grateful to M. (T.) Koch, H. Hollak, S. Kaufmann, C. Fröb, the Light Microscopy Facility and the Fish Facility of the MPI-CBG for experimental help. We thank B. Lombardot and R. Haase from the MPI-CBG Scientific Computing Facility for custom image analysis tools (Script 1 and 2). We also thank the ZIRC for the hdac1 mutant line.

M.M. was a member of the IMPRS of Cell, Developmental and Systems Biology.

The authors declare no competing financial interests.

## Supplementary information

**S1 Figure.**
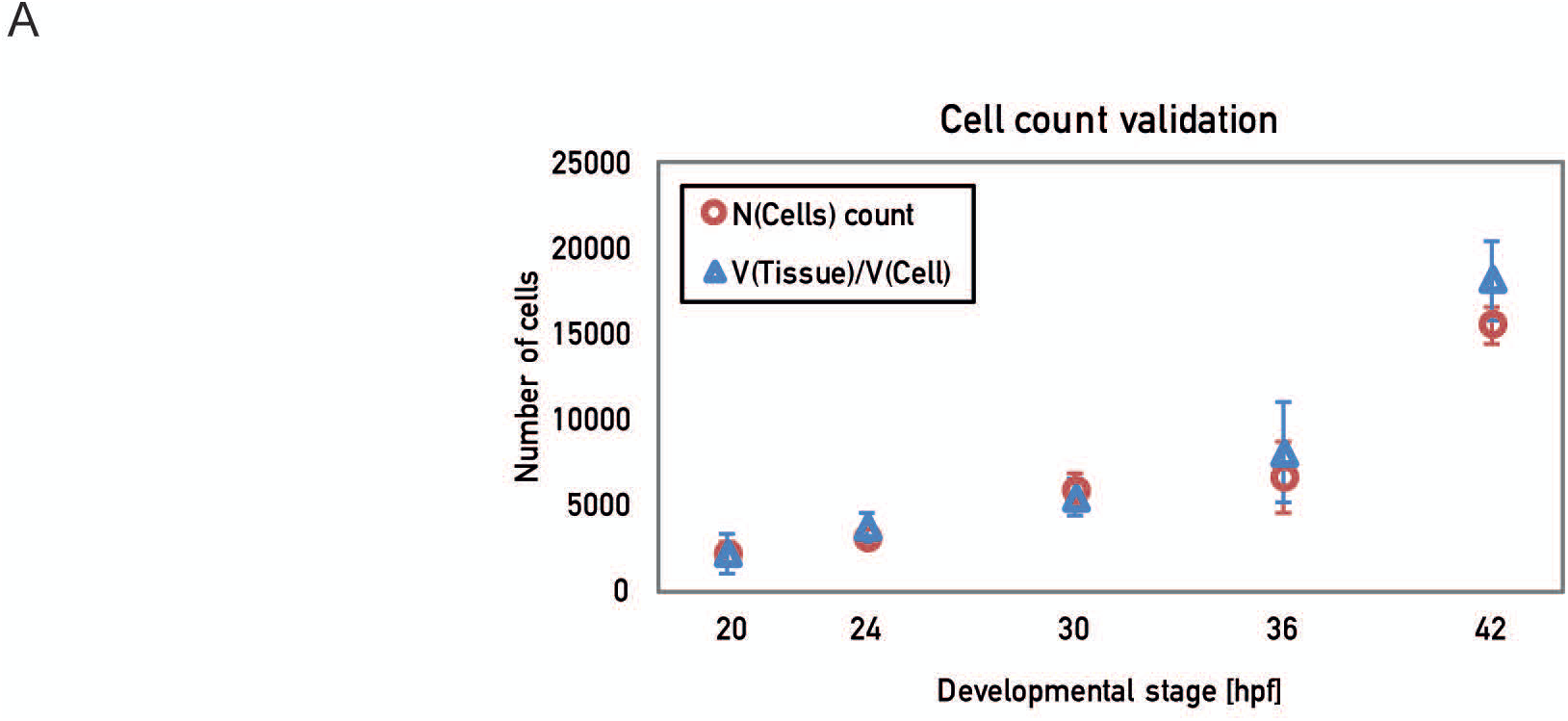
Validation of cell count. [A] The cell count [red] was validated using mitotic cell volumes by dividing the average tissue volume by 90% of mitotic cell average volume for each stage [see Methods]. Data is plotted as stage mean ± S D. n=10 samples/stage for all.

**S2 Figure.**
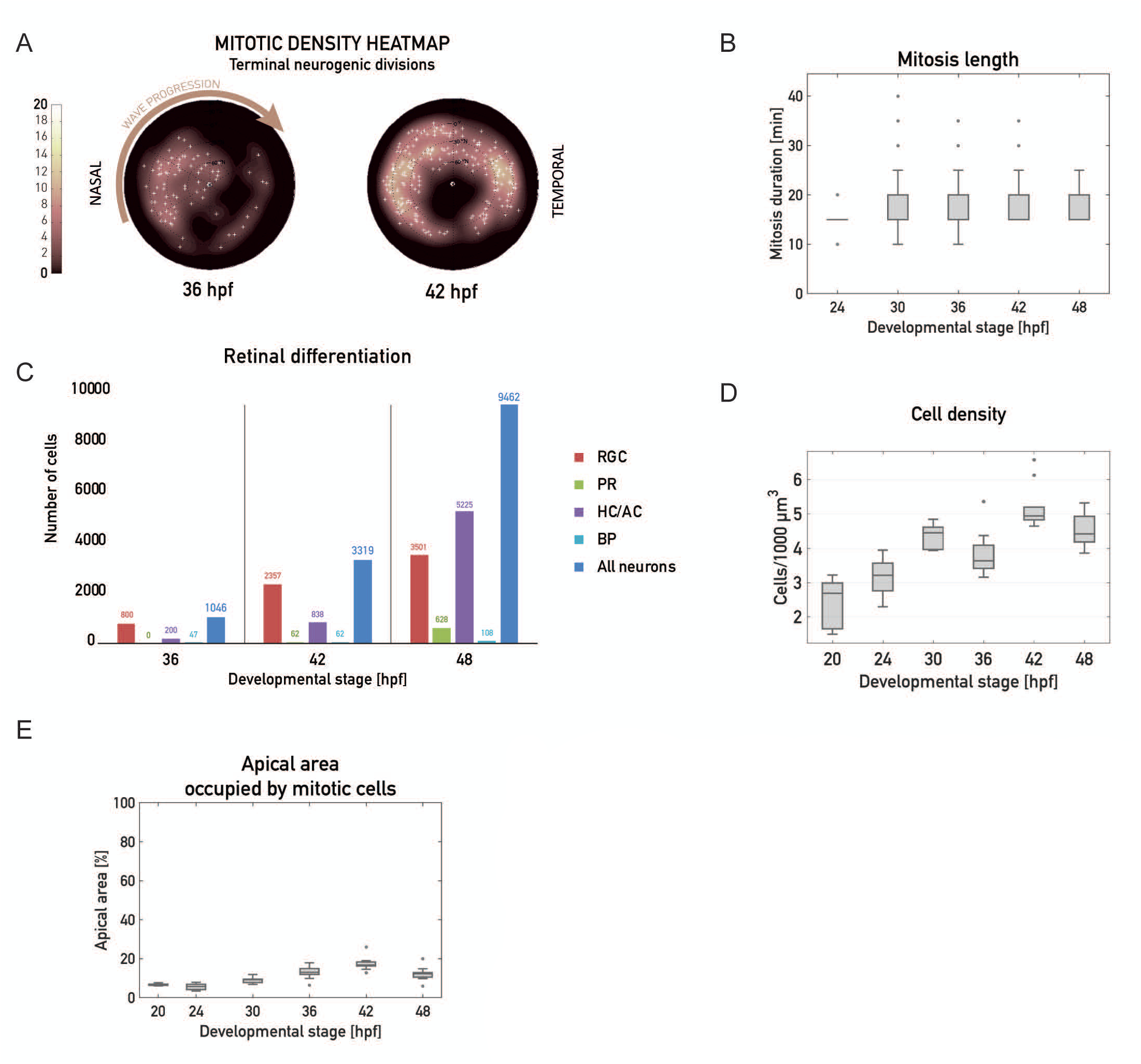
Retinal mitoses and differentiation. **(A)** Heatmaps of Ath5+ [neurogenic) mitotic divisions at 36 hpf and 42 hpf. Validation that non-uniform cell divisions are detectable using the method in Fig. 2c, showing that neurogenic divisions are non-uniform and progress through the tissue as a naso-temporal wave. **(B)** Duration of mitosis does not change over development. Cells labelled mosaically with Hsp70 ::H2B - RFP were tracked in light sheet time-lapses at 5 min time resolution. Data was binned as developmental stage+/-3h. N = 197 cells from 20 embryos [24 hpf N = 20; 30 hpf N = 56; 36 hpf N = 57; 42 hpf N = 53; 48 hpf N=11]. **(C)** Retinal neuro genesis. Average number of neuronal subtypes, as analyzed by FACS sorting from pooled dissected Tg[SoFa) retinal samples. N = 20 retinas/stage. Data was normalized to wildtype background fl uor escence. **(D)** Cell density calculated by dividing the number of cells by total tissue volume. N = 10 samples/stage. **(E)** Fraction of the apical tissue surface area occupied by mitotic cell s. 10 samples/stage.

**S3 Figure.**
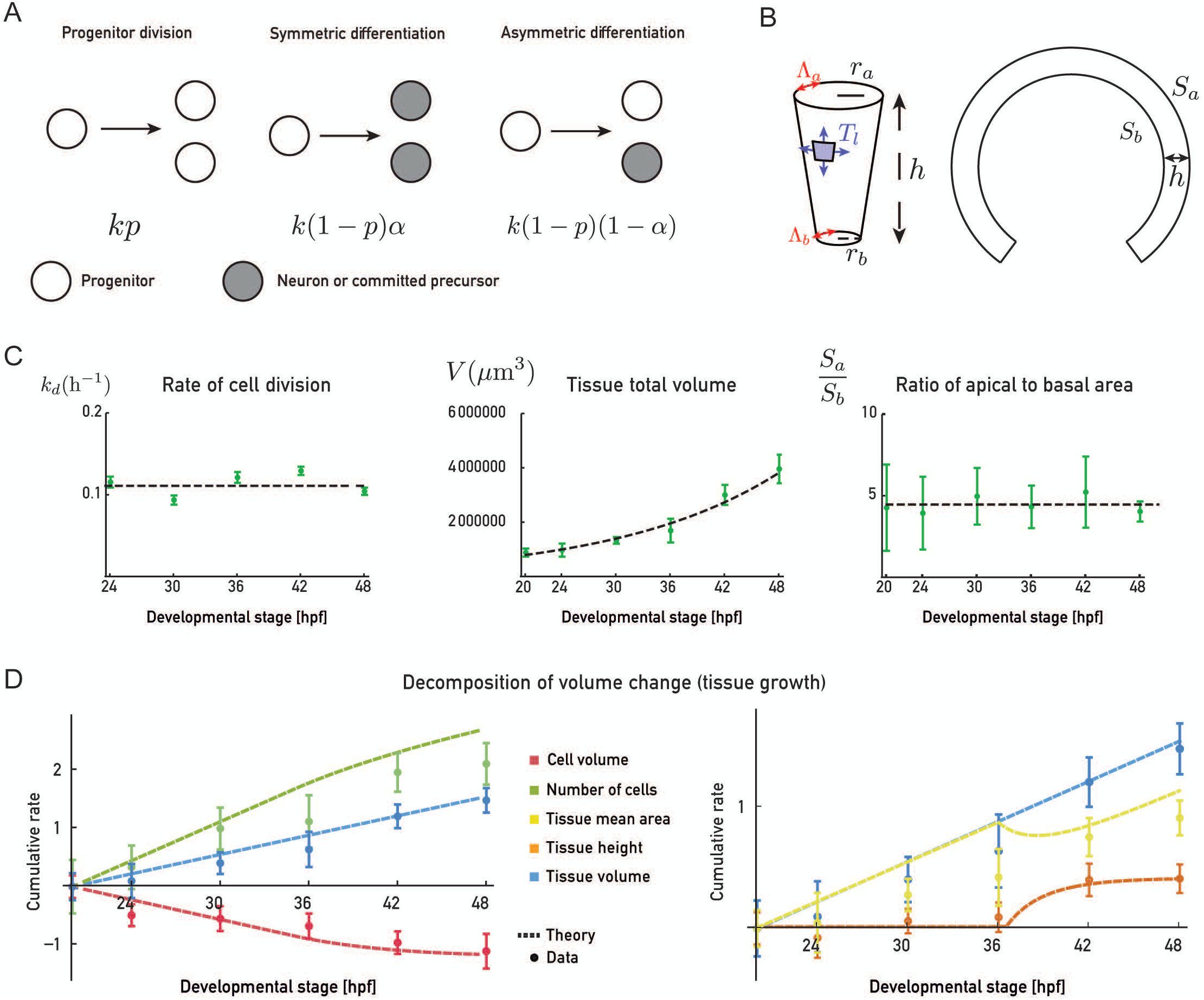
Simplified description of zebrafish retina growth between 20 hpf and 48 hpf. **(A)** Schematic of division and differentiation rules considered in the simplified description of retina growth. For simplicity, we consider 2 cell populations, progenitors [white) and neurons or committed precursors [gray). Progenitors divide with a constant rate ***k.*** Cell division give rise to 2 progenitors with probability ***p,*** 2 neurons/committed precursors with probability **[1-p)a,** one progenitor and one neuron/committed precursor with probability [1-p][1-a]. Here we assume that only symmetric differentiation events occur, such that α=1. **(B)** Schematic of cell and tissue shape geometry. Cells are represented by truncated cones with apical and basal line tensions ***Λ***_***a***_ and ***Λ***_***b***_, and lateral surface tension ***T***_***l***_ The apical and basal tissue surface area is obtained in the simplified description by multiplying the cellular apical and basal area by the number of cells. **(C)** Experimental data for the rate of cell division, tissue volume and ratio of apical to basal surface area as a function of time, and fit to a constant average value [cell division rate and ratio of areas) or to an exponential [tissue volume). **(D)** Comparison between experimentally measured cumulative rates of change of tissue volume, number of cells, cell volume, tissue area and tissue height, and cumulative rates obtained from the simplified description.

**S4 Figure.**
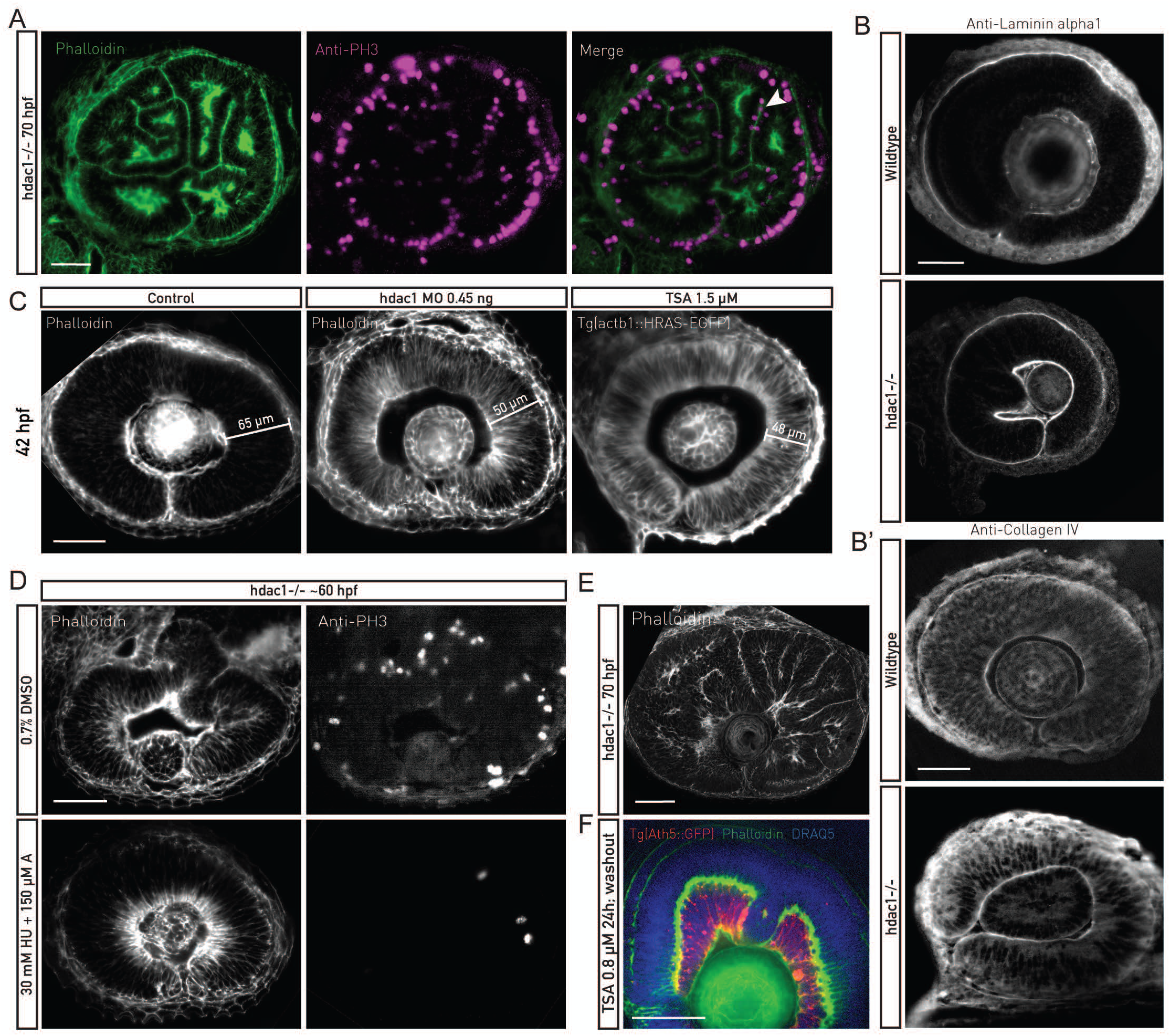
Hdacl -/- characterization. [A] Phalloidin [green] and pH3 antibody [magenta] staining of F-actin and mitotic cells. Apical mitoses, junctional belts and basal actin accumulation are preserved in folded, hdac1-/- retinal PSE. [Bl Laminin-alpha1 and Collagen IV [B“] antibody staining of wildtype and hdac1-/- retinal tissues. The basal lamina underlies the basal surface of the hdac1-/- PSE. [Cl Tissue thickness does not increase in Hdac1 MO-injected or TSA treated retinal PSE [42 hpfl. [DI Phalloidin and pH3 antibody staining of control and HU/A treated hdac1-/-. HU/A treatment started at 42 hpf and lasted for 18 h. HU/A-treated hdac1-/- retinal PSE preserve the basolateral actin accumulation, but do not fold. [El Phalloidin staining of −3 dpf hdac1-/-. Epithelial folds form throughout the retinal PSE. [Fl Tg[At h5::GFP] retinas [red] treated with TSA at 24 hpf for 24 h, and stained with Phalloidin [green] and DRAQ5 [blue]. The medium was replaced after 24 h. The PSE differentiates by 72 hpf despite tissue shape perturbed by folds. Consequently, neuronal layers are pertur bed, as well. Scale bar : 50 µm for all images.

**S5 Figure.**
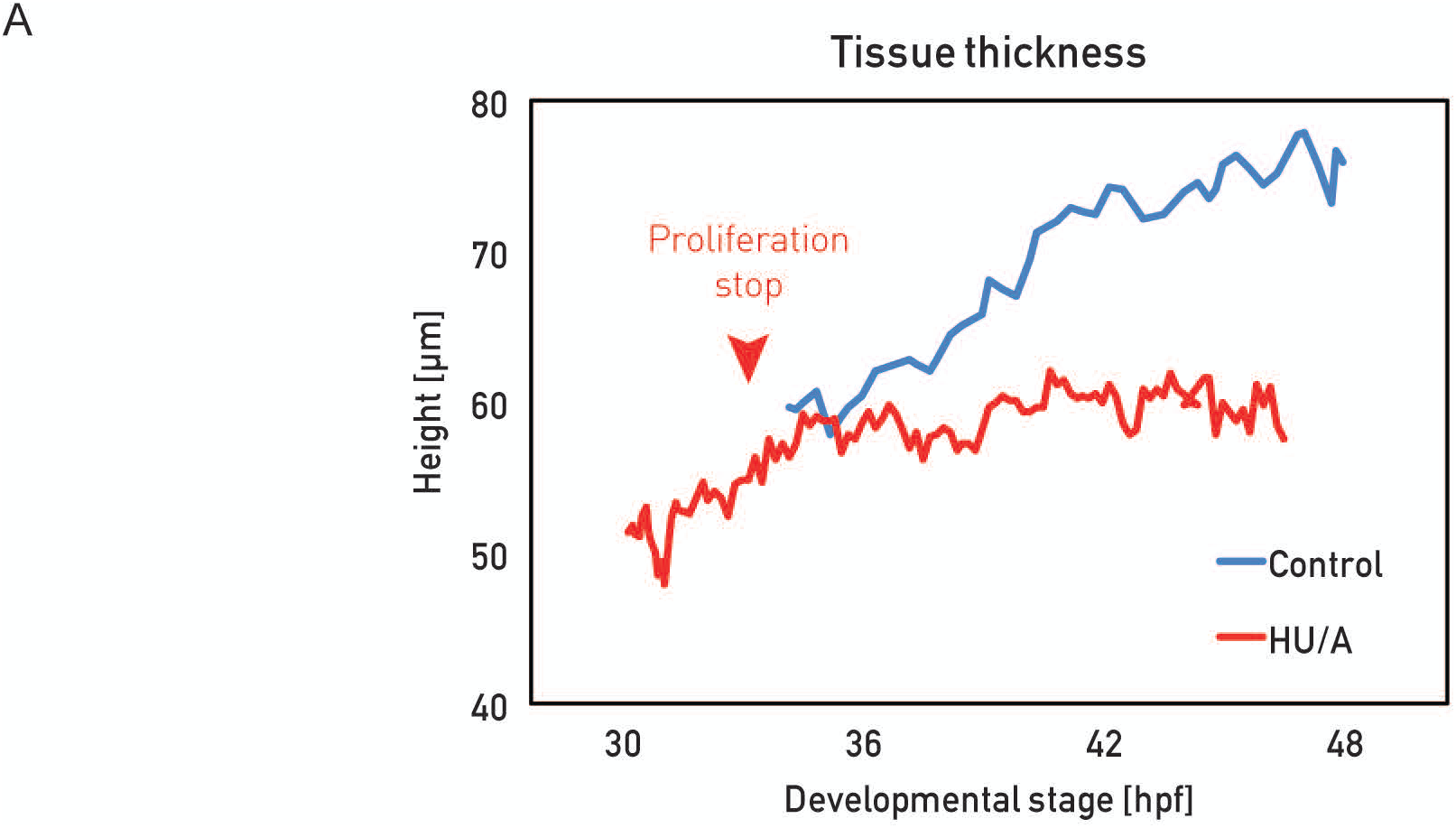
Proliferation is necessary for cell height increase. [A] Cell height from live [light sheet time-lapses] HU/A-treated wildtype retinal PSE. 30 mM HU and 150 µMA were added at the beginning of the movie [30 hpfl. Proliferation stopped 3 h later [red arrowhead]. Cell height did not increase further after cells cycle was blocked.

Movie 1

**3D growth analysis workflow**

Example of retinal neuroepithelial segmentation and analysis of mitotic density. All nuclei are marked with DAPI (blue). Mitotic nuclei are labeled with pH3 antibody (white, later magenta spheres). Spot tool in Imaris was used to automatically detect mitotic nuclei at apical surface (magenta spheres). Surface tool was used in Imaris to manually segment retinal neuroepithelial and lens outlines (grey and light blue).

Scale bar = 50 µm.

Movie 2

**Analysis of cell division angles at apical surface**

Time-lapse Tg(actb1:HRAS-EGFP) (grey) embryos at 35 hpf. View towards apical side of epithelium. Angles of dividing cells were analyzed perpendicular to division plane, using line tool in FIJI.

Time in h:min. Scale bar = 10 µm. Related to Figure 1 and 2 c and d.

Movie 3

**Cell cycle analysis in retinal PSE**

Time-lapse imaging of embryo injected with Hsp70::H2B-RFP (grey) DNA. Red dot marks nucleus followed over multiple cell cycles. Cell cycle length was measured as time between subsequent cell divisions.

Time in h:min. Scale bar = 10 µm. Related to Figure 2g.

Movie 4

**Shape perturbation of hdac1 -/- retinal PSE**

Time-lapse imaging of hdac1 -/- embryo injected with Ras-GFP RNA for uniform labeling (black).

Time in h:min. Scale bar = 50 µm. Related to Figure 5.

Movie 5

**Shape changes over growth in control versus hdac1 -/- retinal PSE**

Time-lapse imaging of control and hdac1 -/- embryos injected with Ras-GFP RNA for uniform labeling (grey).

Time in h:min. Scale bar = 50 µm. Related to Figure 5.

Movie 6

**Tissue wide lateral actin redistribution during retinal PSE growth**

Time-lapse imaging of actin redistribution labelled by Tg(actb1::GFP-UtrophinCH) (heatmap).

Time in h:min. Scale bar = 50 µm. Related to Figure 6.

**Table 1:** Zebrafish transgenic lines and mutant used in this study.

|  | Line/ mutant | Labeling/ purpose | Reference |
| --- | --- | --- | --- |
| 1 | Wild-type WT-AB | Control studies | RRID:ZIRC_ZL1 |
| 2 | Wild-type WT-TL | Control studies | RRID:ZIRC_ZL86 |
| 3 | Tg(actb1:HRAS-EGFP) | All membranes | 1 |
| 4 | Tg(βactin:mKate2-Ras) | All membranes | 2 |
| 5 | Tg(actb1:GFP-UtrCH) | F-actin | 3 |
| 6 | Tg(hsp70:H2B-RFP) | Chromatin | 4 |
| 7 | Tg(ath5:GFP) | Membranes of RGCs and PRs | 2 |
| 8 | Tg(ath5:RFP) | Membranes of RGCs and PRs | 2 |
| 9 | Tg(SoFa2) | Differentiated retinal cells | 5 |
| 10 | hdac1 t24411 | Inhibition of cell cycle exit and differentiation in zebrafish retina | 6 |

**Table 2:**
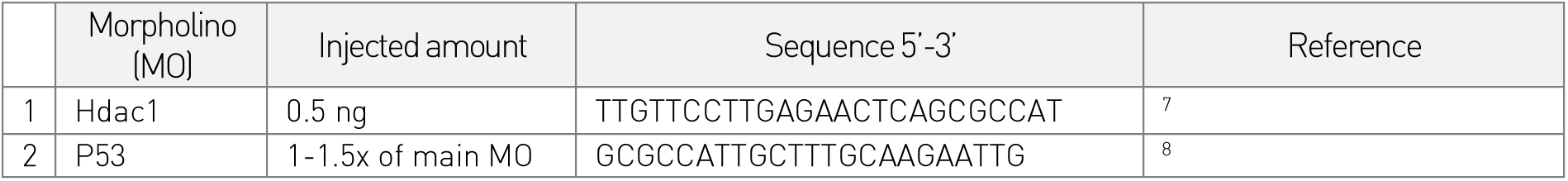
Morpholinos used in this study.

**Table 3:** Chemical inhibitors/drugs used in this study.

|  | Chemical | Function | Working dilution | Source/ Cat.No. |
| --- | --- | --- | --- | --- |
| 1 | Trichostatin A (TSA) | Hdac1 inhibition | 0.8 mM | Sigma-Aldrich; T1952 |
| 2 | Rockout | Rho kinase inhibitor | 175 μM | Santa Cruz; sc-203237 |
| 3 | Hydroxyurea (HU) | S-phase inhibitor | 30 mM | Sigma-Aldrich; H8627-1G |
| 4 | Aphidicolin (A) | S-phase inhibitor | 150 μM | Sigma-Aldrich; 89458 |

## Supplementary Notes

In this supplementary we discuss a set of equations that describe the key aspects of the retina growth between 20 and 48hpf. We take into account balance equations that correspond to the change in number of undifferentiated and neuron or committed precursor cells, and to how the volume growth of the retina is geometrically decomposed in area and height changes. We then introduce a simple description of force balance in the tissue taking into account surface tensions generated along the cellular lateral interfaces, and line tensions generated apically and basally. Taking into account the cell division and differentiation rate, the rate of volume growth, and the change in ratio of lateral surface tension to apicobasal line tension, allows to close the system of equations to obtain a description of the change in time of retina volume, height, area and number of cells.

### I. BALANCE EQUATIONS

We consider two cell types, undifferentiated progenitors (P) and neurons or committed precursors (N). We assume that progenitors divide with a rate *k*. Each division yields two undifferentiated progenitors with probability *p*, two committed neurons with probability (1 − *p*)*α* and one progenitor and one neuron with probability (1 − *p*)(1 − *α*) (Supplementary fig. 3a). Denoting *N*_*p*_ and *N*_*n*_ the number of progenitors and neurons in the retina, and *N*_*t*_ = *N*_*p*_ +*N*_*n*_ the total number of cells, the balance equations for the number of cells then read

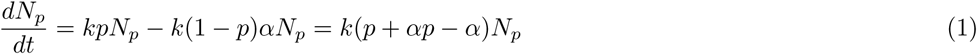

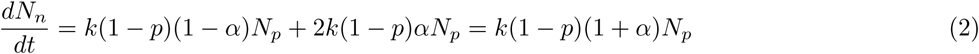

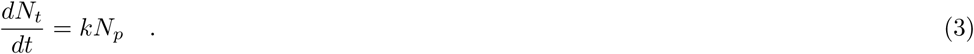

The cell doubling time *T*_2_ is related to the rate of cell division *k* by *T*_2_ = ln 2/*k*. We find that the mean cell cycle length is roughly constant over time, with an average cell doubling time *T*_2_ ≃ 6.2h (Supplementary fig. 3c). This corresponds to an average rate of cell division

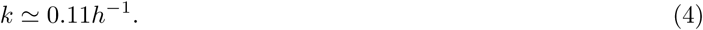

We further make the simplifying approximation here that the number of asymmetric divisions can be neglected, *α* = 1. We also assume that no neuron or committed precursor are present before 35h, as only a low number of differentiated cells can be detected in the retina during this time. Therefore, we assume *p* = 1 prior to 35h. Adjusting for experimental data of the number of differentiated neurons and total number of cells after this time point, we find *p* ≃ 0.65. Specifically, we fitted experimental data of *N*_*t*_(*t*) and *N*_*n*_(*t*) with *N*_*t*_(20hpf) and *p*(*t* > 35hpf) as free parameters. The function *p*(*t*) is then given by:

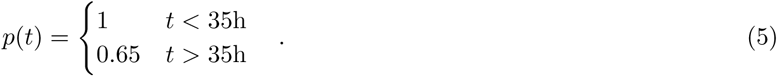

Note that here we chose to do a simple fit to the data taking into account a step function of probability *p*, but the actual value of *p*(*t*) might change in time with a more complicated dynamics [1].

The average cell volume *v* increases due to the overall increase of volume of the tissue, and decreases because of cell division. This can be expressed through the balance equation

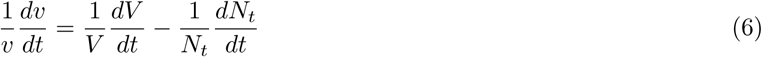

where *V* is the total volume of the retina. We find that the change of total volume is well described by an exponential growth with rate

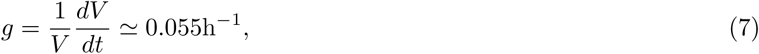

where the rate in Eq. 7 is obtained from a fit of experimental data to the function ln *V* (*t*) = *gt* + *V*_0_, with *g* and *V*_0_ as free parameters (Supplementary Fig. 3c).

Eq. 6 can be integrated to yield

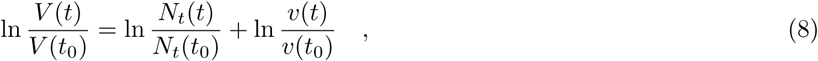

with *t*_0_ = 20hpf. Different terms in Eq. 8 are plotted in Fig. 2b from experimental data to obtain a decomposition of tissue volume growth into average cell volume growth and cell number change. Similarly, in Fig. 2a we plot different terms of the decomposition:

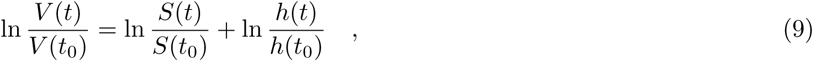

with *S*(*t*) the tissue average surface area and *h*(*t*) the tissue height. Here we make the simplifying assumption that the tissue volume is linearly dependent in the tissue average area and the tissue height.

### II. CELL SHAPE AND MECHANICAL EQUATIONS

We consider here a simple model of cell geometry where cells are described by truncated cones with apical and basal surface area *s*_*a*_ and *s*_*b*_, height *h*, and radius in the middle plane *r* (Supplementary Fig. 3b). The middle plane is defined here as the plane with cross-sectional surface area equal to the mean apical and basal surface areas. We do not discuss the origin of cell and tissue curvature, which is captured in a relative area parameter *s* = *s*_*a*_/*s*_*b*_, taken to be constant and equal on average to ~ 4.5 from experimental data (Supplementary Fig. 3c). The cell apical area is given by 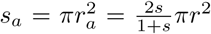 and the basal area is given by 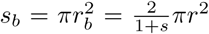. As a result, the radii of the apical and basal cell surfaces are given by 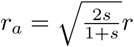 and 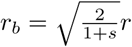, the apical and basal perimeters by *p*_*a*_ = 2π*r*_*a*_ and *p*_*b*_ = 2π*r*_*b*_, the lateral surface area by 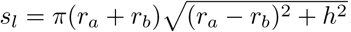. The cell volume *v* is given by

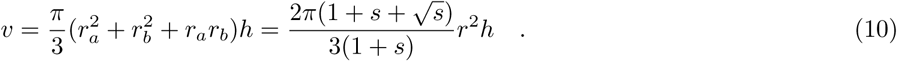

To obtain the total average tissue area, we consider that the retina is made of identical cells, such that:

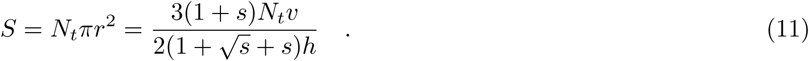

Deviations from this ideal behaviour, for instance due to the fact that cells of the retina margin have different shapes than cells in the central part of the retina, may be at the origin of differences between the area predicted by the simplified model and experimentally measured areas (Fig. 4g and Supplementary Fig. 3d).

To describe cell mechanics, we consider a simple model where cells have line tensions around the apical and basal perimeter Λ_*a*_ and Λ_*b*_, and surface tension along their lateral surface *T*_*l*_/2. The line tensions Λ_*a*_ can arise from the apical actomyosin epithelial belt, the line tensions Λ_*b*_ from the basal actomyosin belt. The surface tension *T*_*l*_ can arise from the actomyosin cortex exerting tension along cellular interfaces. The mechanical energy for one cell is then written

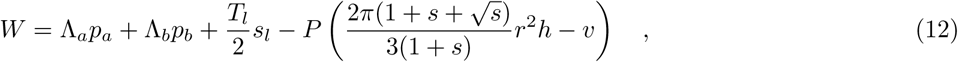

where geometrical quantities have been introduced above, and *P* is a Lagrange multiplier enforcing the volume constraint and corresponding to the intracellular pressure. Minimizing *W* with respect to *r*, *h* and *P*, one obtains the equilibrium cell height written here in the limit of highly columnar cells, *h ≫ r*:

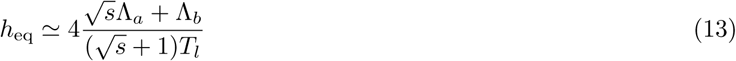

We further assume that dissipative contributions slow down the height relaxation on a time scale *τ*_*h*_, such that

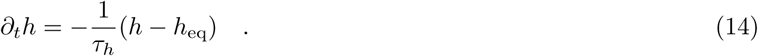

Therefore, in this simple model, the cell height is set by the ratio 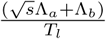. Actin intensity measurements (Fig. 4b) indicate that the ratio of apical and basal actin to lateral actin concentration increases weakly before 36hAPF and more strongly at 42hAPF. We assume here that observed changes in these actin ratio lead to a change in the ratio of apicobasal line tensions and lateral surface tensions, such that:

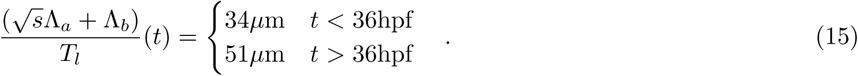

where this value were chosen to match the experimentally measured cell height change (Fig. 4e).

### III. TISSUE GROWTH DYNAMICS

Combining the balance equations 1-7 and the force balance equations 13-14 with apico-basal line tension to lateral surface tension ratio given by eq. 15, we can solve for the evolution of time of the parameters characterizing tissue growth. We plot in Fig. 2h, Fig. 4e-4g and Supplementary Fig. 3d the resulting curves for the total number of cells and the number of neurons, the cell volume, the tissue height and the tissue area. Parameters are determined from fits to experimental data, except for *τ*_*h*_ which we take equal to 3h, as the data time resolution does not allow for a more precise determination. The agreement between the simplified theory and experimental data corresponds to the following picture of retinal growth:

- Tissue volume accumulation and progenitor cell division occur at a roughly constant rate.
- Before 35h, there is no cell differentiation, and the actomyosin distribution is maintained, such that forces generated by the actomyosin cytoskeleton ensure a constant tissue height. Because the rate of volume growth *g* ≃ 0.055h^−1^, is about twice lower than the rate of cell division *k* ≃ 0.11*h*^−1^, the cell volume is decreasing (Fig. 4c).
- After 35h, ~35% of cell divisions lead to cell differentiation. At the same time, the actomyosin distribution is changing, leading to cell elongation. The relative rate of new cell production slows down, due to terminally differentiated cells stopping division. As a result, the relative rate of cell volume decrease is also reduced.

In Fig. 5g we plot the tissue aspect ratio calculated from the equations described in section II, as well as for a modified model where the ratio 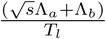 does not change and is equal to 34*µ*m. This corresponds to a situation where the actomyosin distribution does not change, and as a result the cell height is constant. In that case the tissue aspect ratio keeps decreasing after 36hpf (Fig. 5g).

## Footnotes

[1] Jie He, Gen Zhang, Alexandra D Almeida, Michel Cayouette, Benjamin D Simons, and William A Harris. How variable clones build an invariant retina. *Neuron*, 75(5):786–798, 2012.

## Abbreviations

CoV: Coefficient of Variation
ECM: extracellular matrix
hdac1: histone-deacetylase 1
hpf: hours post fertilization
pH3: phosphorylated histone H3
PSE: pseudostratified epithelium

## References

1. Andrew DJ, Henderson KD, Seshaiah P. Salivary gland development in Drosophila melanogaster. Mechanisms of Development. Elsevier; 2000;92: 5–17. doi:10.1016/S0925-4773(99)00321-4

2. Sidhaye J, Norden C. Concerted action of neuroepithelial basal shrinkage and active epithelial migration ensures efficient optic cup morphogenesis. eLife. eLife Sciences Publications Limited; 2017;6: 73. doi:10.7554/eLife.22689

3. Behrndt M, Salbreux G, Campinho P, Hauschild R, Oswald F, Roensch J, et al. Forces driving epithelial spreading in zebrafish gastrulation. Science. 2012;338: 257–260. doi:10.1126/science.1224143

4. Collinet C, Rauzi M, Lenne P-F, Lecuit T. Local and tissue-scale forces drive oriented junction growth during tissue extension. Nature Cell Biology. Nature Publishing Group; 2015;17: ncb3226–1258. doi:10.1038/ncb3226

5. Rauzi M, Krzic U, Saunders TE, Krajnc M, Ziherl P, Hufnagel L, et al. Embryo-scale tissue mechanics during Drosophila gastrulation movements. Nature Communications. 2015;6: 8677. doi:10.1038/ncomms9677

6. He L, Wang X, Tang HL, Montell DJ. Tissue elongation requires oscillating contractions of a basal actomyosin network. Nature Cell Biology. Nature Publishing Group; 2010;12: 1133–1142. doi:10.1038/ncb2124

7. Martin AC, Gelbart M, Fernandez-Gonzalez R, Kaschube M, Wieschaus EF. Integration of contractile forces during tissue invagination. J Cell Biol. Rockefeller University Press; 2010;188: 735–749. doi:10.1083/jcb.200910099

8. Galea GL, Cho Y-J, Galea G, Molè MA, Rolo A, Savery D, et al. Biomechanical coupling facilitates spinal neural tube closure in mouse embryos. Proc Natl Acad Sci USA. National Academy of Sciences; 2017;114: E5177–E5186. doi:10.1073/pnas.1700934114

9. Booth AJR, Blanchard GB, Adams RJ, Röper K. A dynamic microtubule cytoskeleton directs medial actomyosin function during tube formation. Developmental Cell. 2014;29: 562–576. doi:10.1016/j.devcel.2014.03.023

10. Grego-Bessa J, Bloomekatz J, Castel P, Omelchenko T, Baselga J, Anderson KV. The tumor suppressor PTEN and the PDK1 kinase regulate formation of the columnar neural epithelium. eLife. eLife Sciences Publications, Ltd; 2016;5: 20806. doi:10.7554/eLife.12034

11. Etournay R, Popović M, Merkel M, Nandi A, Blasse C, Aigouy B, et al. Interplay of cell dynamics and epithelial tension during morphogenesis of the Drosophila pupal wing. eLife. eLife Sciences Publications Limited; 2015;4: e07090. doi:10.7554/eLife.07090

12. Dye NA, Popović M, Spannl S, Etournay R, Kainmüller D, Ghosh S, et al. Cell dynamics underlying oriented growth of the Drosophila wing imaginal disc. Development. Oxford University Press for The Company of Biologists Limited; 2017;144: dev.155069–4421. doi:10.1242/dev.155069

13. Bardet P-L, Guirao B, Paoletti C, Serman F, Léopold V, Bosveld F, et al. PTEN Controls Junction Lengthening and Stability during Cell Rearrangement in Epithelial Tissue. Developmental Cell. Elsevier; 2013;25: 534–546. doi:10.1016/j.devcel.2013.04.020

14. Fuhrmann S. Eye Morphogenesis and Patterning of the Optic Vesicle. Current topics in developmental biology. NIH Public Access; 2010;93: 61–84. doi:10.1016/B978-0-12-385044-7.00003-5

15. Strzyz PJ, Matejcic M, Norden C. Heterogeneity, Cell Biology and Tissue Mechanics of Pseudostratified Epithelia: Coordination of Cell Divisions and Growth in Tightly Packed Tissues. Int Rev Cell Mol Biol. Elsevier; 2016;325: 89–118. doi:10.1016/bs.ircmb.2016.02.004

16. Strzyz PJ, Lee HO, Sidhaye J, Weber IP, Leung LC, Norden C. Interkinetic nuclear migration is centrosome independent and ensures apical cell division to maintain tissue integrity. Developmental Cell. 2015;32: 203–219. doi:10.1016/j.devcel.2014.12.001

17. Smart IH. Proliferative characteristics of the ependymal layer during the early development of the spinal cord in the mouse. J Anat. Wiley-Blackwell; 1972;111: 365–380. doi:10.1111/(ISSN)1469-7580

18. Norden C. Pseudostratified epithelia - cell biology, diversity and roles in organ formation at a glance. J Cell Sci. The Company of Biologists Ltd; 2017;130: 1859–1863. doi:10.1242/jcs.192997

19. Miyata T, Okamoto M, Shinoda T, Kawaguchi A. Interkinetic nuclear migration generates and opposes ventricular-zone crowding: insight into tissue mechanics. Front Cell Neurosci. Frontiers; 2015;8: 673. doi:10.3389/fncel.2014.00473

20. Dzafic E, Strzyz PJ, Wilsch-Bräuninger M, Norden C. Centriole Amplification in Zebrafish Affects Proliferation and Survival but Not Differentiation of Neural Progenitor Cells. Cell Rep. 2015;13: 168–182. doi:10.1016/j.celrep.2015.08.062

21. Icha J, Weber M, Waters JC, Norden C. Phototoxicity in live fluorescence microscopy, and how to avoid it. BioEssays. 2017;39. doi:10.1002/bies.201700003

22. He J, Zhang G, Almeida AD, Cayouette M, Simons BD, Harris WA. How variable clones build an invariant retina. Neuron. 2012;75: 786–798. doi:10.1016/j.neuron.2012.06.033

23. Almeida AD, Boije H, Chow RW, He J, Tham J, Suzuki SC, et al. Spectrum of Fates: a new approach to the study of the developing zebrafish retina. Development. 2014;141: 1971–1980. doi:10.1242/dev.104760

24. Weber IP, Ramos AP, Strzyz PJ, Leung LC, Young S, Norden C. Mitotic position and morphology of committed precursor cells in the zebrafish retina adapt to architectural changes upon tissue maturation. Cell Rep. 2014;7: 386–397. doi:10.1016/j.celrep.2014.03.014

25. Fish JL, Dehay C, Kennedy H, Huttner WB. Making bigger brains-the evolution of neural-progenitor-cell division. J Cell Sci. The Company of Biologists Ltd; 2008;121: 2783–2793. doi:10.1242/jcs.023465

26. Okamoto M, Namba T, Shinoda T, Kondo T, Watanabe T, Inoue Y, et al. TAG1–assisted progenitor elongation streamlines nuclear migration to optimize subapical crowding. Nature Neuroscience. 2013;16: 1556–1566. doi:10.1038/nn.3525

27. Salbreux G, Charras G, Paluch E. Actin cortex mechanics and cellular morphogenesis. Trends in Cell Biology. 2012;22: 536–545.

28. Lecuit T, Lenne P-F. Cell surface mechanics and the control of cell shape, tissue patterns and morphogenesis. Nature Reviews Molecular Cell Biology. 2007;8: 633–644. doi:10.1038/nrm2222

29. Paluch E, Heisenberg C-P. Biology and Physics of Cell Shape Changes in Development. Current Biology. Elsevier; 2009;19: R790–R799. doi:10.1016/j.cub.2009.07.029

30. Yamaguchi M, Tonou-Fujimori N, Komori A, Maeda R, Nojima Y, Li H, et al. Histone deacetylase 1 regulates retinal neurogenesis in zebrafish by suppressing Wnt and Notch signaling pathways. Development. The Company of Biologists Ltd; 2005;132: 3027–3043. doi:10.1242/dev.01881

31. Stadler JA, Shkumatava A, Norton WHJ, Rau MJ, Geisler R, Fischer S, et al. Histone deacetylase 1 is required for cell cycle exit and differentiation in the zebrafish retina. Dev Dyn. Wiley-Liss, Inc; 2005;233: 883–889. doi:10.1002/dvdy.20427

32. Ma M, Cao X, Dai J, Pastor-Pareja JC. Basement Membrane Manipulation in Drosophila Wing Discs Affects Dpp Retention but Not Growth Mechanoregulation. Developmental Cell. 2017;42: 97–106. doi:10.1016/j.devcel.2017.06.004

33. Harunaga JS, Doyle AD, Yamada KM. Local and global dynamics of the basement membrane during branching morphogenesis require protease activity and actomyosin contractility. Developmental Biology. 2014;394: 197–205. doi:10.1016/j.ydbio.2014.08.014

34. Kwan KM. Coming into focus: the role of extracellular matrix in vertebrate optic cup morphogenesis. Mansour SL, Ornitz DM, editors. Developmental Dynamics. 2014;243: 1242–1248. doi:10.1002/dvdy.24162

35. Elosegui-Artola A, Jorge-Peñas A, Moreno-Arotzena O, Oregi A, Lasa M, García-Aznar JM, et al. Image Analysis for the Quantitative Comparison of Stress Fibers and Focal Adhesions. Janody F, editor. PLoS ONE. Public Library of Science; 2014;9: e107393. doi:10.1371/journal.pone.0107393

36. Calderwood DA, Shattil SJ, Ginsberg MH. Integrins and actin filaments: reciprocal regulation of cell adhesion and signaling. J Biol Chem. American Society for Biochemistry and Molecular Biology; 2000;275: 22607–22610. doi:10.1074/jbc.R900037199

37. Hodkinson PS, Elliott PA, Lad Y, McHugh BJ, MacKinnon AC, Haslett C, et al. Mammalian NOTCH-1 activates beta1 integrins via the small GTPase R-Ras. J Biol Chem. American Society for Biochemistry and Molecular Biology; 2007;282: 28991–29001. doi:10.1074/jbc.M703601200

38. Leong KG, Hu X, Li L, Noseda M, Larrivée B, Hull C, et al. Activated Notch4 inhibits angiogenesis: role of beta 1-integrin activation. Mol Cell Biol. 2002;22: 2830–2841.

39. Gutzman JH, Graeden E, Brachmann I, Yamazoe S, Chen JK, Sive H. Basal constriction during midbrain-hindbrain boundary morphogenesis is mediated by Wnt5b and Focal Adhesion Kinase. bioRxiv. Cold Spring Harbor Laboratory; 2018; 251132. doi:10.1101/251132

40. Shraiman BI. Mechanical feedback as a possible regulator of tissue growth. Proc Natl Acad Sci USA. National Acad Sciences; 2005;102: 3318–3323. doi:10.1073/pnas.0404782102

41. Pantazis P, Supatto W. Advances in whole-embryo imaging: a quantitative transition is underway. Nature Reviews Molecular Cell Biology. 2014;15: 327–339. doi:10.1038/nrm3786

42. Paşca SP. The rise of three-dimensional human brain cultures. Nature. Nature Publishing Group; 2018;553: 437. doi:10.1038/nature25032

43. Zhong X, Gutierrez C, Xue T, Hampton C, Vergara MN, Cao L-H, et al. Generation of three-dimensional retinal tissue with functional photoreceptors from human iPSCs. Nature Communications. 2014;5: 147. doi:10.1038/ncomms5047

44. Llonch S, Carido M, Ader M. Organoid technology for retinal repair. Developmental Biology. Academic Press; 2018;433: 132–143. doi:10.1016/j.ydbio.2017.09.028

45. Kimmel CB, Ballard WW, Kimmel SR, Ullmann B, Schilling TF. Stages of embryonic development of the zebrafish. Developmental Dynamics. Wiley Subscription Services, Inc., A Wiley Company; 1995;203: 253–310. doi:10.1002/aja.1002030302

46. Inoue D, Wittbrodt J. One for All—A Highly Efficient and Versatile Method for Fluorescent Immunostaining in Fish Embryos. Callaerts P, editor. PLoS ONE. Public Library of Science; 2011;6: e19713. doi:10.1371/journal.pone.0019713

47. Schindelin J, Arganda-Carreras I, Frise E, Kaynig V, Longair M, Pietzsch T, et al. Fiji: an open-source platform for biological-image analysis. Nat Methods. 2012;9: 676–682. doi:10.1038/nmeth.2019

## SUPPLEMENTARY REFERENCES

1. Cooper, M. S. et al. Visualizing morphogenesis in transgenic zebrafish embryos using BODIPY TR methyl ester dye as a vital counterstain for GFP. Dev. Dyn. 232, 359–368 (2005).

2. Icha, J., Kunath, C., Rocha-Martins, M. & Norden, C. Independent modes of ganglion cell translocation ensure correct lamination of the zebrafish retina. The Journal of Cell Biology 215, 259–275 (2016).

3. Behrndt, M. et al. Forces driving epithelial spreading in zebrafish gastrulation. Science 338, 257–260 (2012).

4. Dzafic, E., Strzyz, P. J., Wilsch-Bräuninger, M. & Norden, C. Centriole Amplification in Zebrafish Affects Proliferation and Survival but Not Differentiation of Neural Progenitor Cells. Cell Rep 13, 168–182 (2015).

5. Almeida, A. D. et al. Spectrum of Fates: a new approach to the study of the developing zebrafish retina. Development 141, 1971–1980 (2014).

6. Stadler, J. A. et al. Histone deacetylase 1 is required for cell cycle exit and differentiation in the zebrafish retina. Dev. Dyn. 233, 883–889 (2005).

7. Yamaguchi, M. et al. Histone deacetylase 1 regulates retinal neurogenesis in zebrafish by suppressing Wnt and Notch signaling pathways. Development 132, 3027–3043 (2005).

8. Robu, M. E. et al. p53 Activation by Knockdown Technologies. PLOS Genet 3, e78 (2007).

